# Pharmaco-genetic evidence of presymptomatic lysosomal dysfunction coupled to detrimental DPR secretion in *Drosophila* Models of C9orf72-linked ALS/FTD

**DOI:** 10.64898/2026.11.09.765393

**Authors:** Paulien H Smeele, Veronica Ferrari, Giorgia Spina, Erika Donà, Ruben Cauchi, Angelo Poletti, Thomas Vaccari

**Author notes:** Department of Physiology and Biochemistry, Faculty of Medicine and Surgery, University of Malta, Msida, Malta (Paulien H Smeele).

## Abstract

Hexanucleotide repeat expansions in *C9orf72*, the most common genetic cause of Amyotrophic lateral sclerosis (ALS) and frontotemporal dementia (FTD), drive neurodegeneration through both loss of gene function and toxic gain-of-function mechanisms, including transcription of repeat RNA and repeat-associated non-AUG (RAN) translation into dipeptide repeat proteins (DPRs). Both disease mechanisms have been linked to disruption of the autophagy-lysosome pathway (ALP), though how this contributes to disease progression remains unclear. By expressing toxic *C9orf72* repeats in *Drosophila melanogaster*, we observed presymptomatic lysosome impairment, coupled with largely preserved autophagosome formation. Pharmacological treatment with valproic acid (VPA) partially prevented neurodegenerative and locomotor phenotypes in expressing animals in presence of Mitf, which is required for lysosomal biogenesis. Genetic inhibition of residual lysosomal degradation enhanced neurotoxicity, whereas downregulation of autophagy and secretory autophagy genes suppressed toxicity, uncovering a previously unrecognized contribution of secretion to disease. Consistent with this, *in vitro* experiments indicated that disabling lysosome function, but not autophagy induction, leads to an increase in neuronal release of autophagy-associated proteins and DPRs, which can be found be uptaken by nearby neurons and glia *in vivo*. We propose that repeat-driven presymptomatic lysosomal dysfunction, prevented by pharmacologic treatment, promotes secretory autophagy, which, in turn facilitates extracellular release and intercellular propagation of neurotoxic DPRs.

## Introduction

Amyotrophic lateral sclerosis (ALS) and frontotemporal dementia (FTD) are fatal neurodegenerative disorders with overlapping clinical and pathological features **(Ling et al., 2013; Masrori & Van Damme, 2020)**. The most common genetic cause of both is a GGGGCC (G4C2) hexanucleotide repeat expansion in the *C9orf72* gene **(DeJesus-Hernandez et al., 2011; Renton et al., 2011)**. The expansion contributes to disease through two mechanisms: loss of C9orf72 function, and/or toxic gain of function mediated by repeat RNA and dipeptide repeat proteins (DPRs) Poly-GA, -GP, -GR, -PA, -PR generated through Repeat-Associated Non-AUG (RAN) translation **(Gendron et al., 2013; Mori et al., 2013; Zu et al., 2013)**.

DPR inclusions are a defining neuropathological feature of C9orf72-linked ALS/FTD and co-localize with p62 and ubiquitin in patient brain tissue **(Mackenzie et al., 2015; May et al., 2014; Ramos-Campoy et al., 2018)**, implicating proteostasis pathways, including the ubiquitin proteosome system and the autophagy-lysosome pathway (ALP), in disease. Consistent with this, ALP dysfunction has been reported across several C9orf72 models. Loss of C9orf72 function appears to impair autophagy initiation and lysosome biogenesis through disrupted ULK1 complex recruitment, Rab1a regulation, and lysosomal maturation **(Amick et al., 2016; Beckers et al., 2023; Corbier & Sellier, 2017; O’Rourke et al., 2016; Sullivan et al., 2016; Webster et al., 2016; Yang et al., 2016)**. In parallel, gain-of-function mechanisms driven by G4C2 repeat RNA and DPRs have been shown to disrupt autophagosome (AP) formation, reduce cargo turnover, and impair lysosomal acidification in *Drosophila* neurons and mammalian cells **(Boivin et al., 2020; Božič et al., 2022; Cunningham et al., 2020; Sung & Lloyd, 2023; Wang et al., 2023)**. Together, these findings indicate that ALP defects arise through both loss- and gain-of-function pathways in C9orf72-linked ALS/FTD.

A growing body of work suggests that these ALP defects may associate with impaired activity of TFEB, the master transcriptional regulator of genes controlling ALP. Studies in *Drosophila* and cultured neurons show that G4C2 repeat expression reduces TFEB (Mitf in *Drosophila)* nuclear import **(Cunningham et al., 2020; Lin et al., 2024; Wang et al., 2023)**, consistent with broader nucleo-cytoplasmic transport impairment **(Freibaum et al., 2015; Zhang et al., 2015)**. However, despite growing evidence linking impaired TFEB activity to C9orf72-linked ALS/FTD pathogenesis, it remains unclear how early these defects arise *in vivo*, how they contribute to disease progression and whether restoring TFEB activity can modify disease phenotypes.

While the ALP is primarily considered a degradative pathway, autophagy-associated vesicles can also be redirected away from lysosomal degradation towards extracellular release. This process, termed autophagy-dependent secretion or secretory autophagy, mediates the unconventional extracellular release of diverse cytosolic cargoes, including cytokines **(Dupont et al., 2011; Kimura et al., 2017)** and aggregation-prone proteins associated with neurodegenerative disease **(Ejlerskov et al., 2013; Nakamura et al., 2024)**. Recent work suggests that secretory autophagy becomes engaged under cellular stress and lysosomal dysfunction **(Solvik et al., 2022)**, and in some contexts can drive neuroinflammation and neurodegeneration **(Hartmann et al., 2024)**.

In mammals, autophagy proteins including Atg5, Atg7 and Atg8/LC3, together with the SNARE proteins Sec22b, Snap29, and Rab8a, act as key regulators of secretory autophagy **(Kimura 2017; Hartmann et al., 2024)**. These mechanisms are conserved in *Drosophila*, in which Snap29, Rab8, and Sec22 have been shown to regulate autophagy-dependent secretion. Notably, Snap29 plays a conserved role in both AP-lysosome fusion and AP-plasma membrane fusion, potentially acting to control either vesicle fate **(Chang et al., 2024).** Thus, the secretory branch of the ALP may become engaged when lysosomal degradative capacity is limited.

In the context of C9orf72-linked ALS/FTD, DPRs are known to propagate between cells in culture **(Chang et al., 2016; Khosravi et al., 2020; Westergard et al., 2016; Zhou et al., 2017)**, and polyGP is detectable in cerebrospinal fluid of patients **(Lehmer et al., 2017; Su et al., 2014)**, indicating the presence of extracellular DPRs *in vivo*. Recent work in *Drosophila* has further shown that polyGA can spread between neurons within the adult brain **(Morón-Oset et al., 2019)**. However, whether secretory autophagy may contribute to DPR spreading has not been explored.

Here, we use complementary *Drosophila* and motor neuron cell models to define how disruption of the ALP pathway contributes to C9orf72 repeat toxicity. We show that Mitf-dependent lysosomal gene expression is selectively suppressed before overt behavioral decline, accompanied by early accumulation of ALP cargoes and expansion of the lysosomal compartments. Through genetic dissection, we identify opposing roles for lysosomal degradation and upstream autophagy in modulating toxicity, and implicate components of the secretory autophagy pathway. We further show that lysosomal impairment is associated with extracellular release of polyGA and autophagy-associated proteins *in vitro* and we study widespread propagation of polyGR in the adult fly brain *in vivo.* Finally, we demonstrate pharmacological Mitf-dependent prevention of G4C2-induced locomotor dysfunction *in vivo*. Together, our findings identify presymptomatic lysosomal dysfunction as a key pathogenic event in a model of C9orf72-linked ALS/FTD and suggest that altered handling of autophagy-associated cargo may contribute to the spread of DPRs and to toxicity.

## Results

### Lysosomal dysfunction precedes neurodegeneration in Drosophila ALS/FT models

We investigated disease progression using established *Drosophila* models expressing either 3 G4C2 repeats (3R; non-pathogenic control), or 36 G4C2 repeats (36R) known to form RNA foci and to undergo RAN translation to generate the sense-derived DPR polyGA, polyGP and polyGR **(Mizielinska et al., 2014).** Under the control of *D42-GAL4,* a driver expressed in the central brain (CB) neurons and in motor neurons of the ventral nerve cord (VNC; **Figure S1A**), 36R expression resulted in normal locomotor function at 7 days post-eclosion (dpe), whereas by 13 dpe flies exhibited a severe locomotor deficit **(Figure 1A)**, consistent with the progressive nature of C9orf72-linked ALS/FTD in patients. We therefore refer to these timepoints as “presymptomatic” and “symptomatic”, respectively.

**Figure 1.**
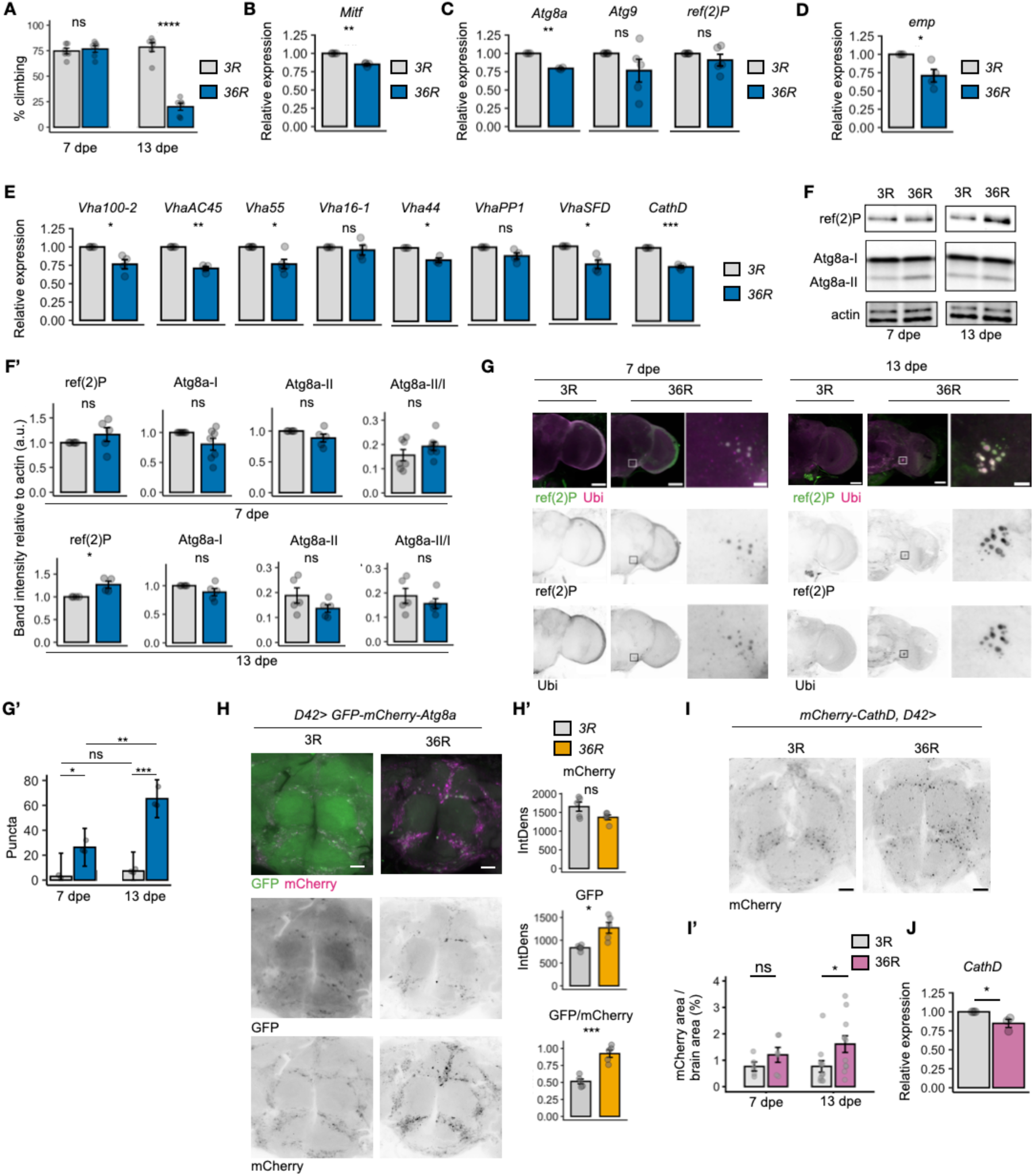
Early ALP disfunction in 36R-expressing flies. **(A)** Percentage of normally climbing flies when expressing 3R or 36R G4C2 repeats in D42-positive neurons at 7 and 13 dpe. Data are shown as mean ± SEM. Unpaired Welch’s t-test; n = 6 biological replicates. **(B-E)** qPCR on adult heads at 7 dpe for *Mitf* (B), autophagy-related genes (*Atg8a*, *Atg9*, *ref(2)P)* (C), *emp* (D), and lysosomal/v-ATPase genes (*Vha100*-2, *VhaAC45*, *Vha55*, *Vha16*-1, *Vha44*, *VhaPP1*, *VhaSFD*, and *CathD*) (E). Gene expression is shown as mean fold change (2^−ΔΔCt^) relative to control. Data are shown as mean ± SEM. Paired Student’s t-tests performed on ΔCt values; n ≥ 3 biological replicates. **(F-F’)** Western blot of ref(2)P, Atg8a and actin in adult heads at 7 and 13 dpe (F) and quantification of band intensity relative to actin (F’). Data are shown as mean ± SEM. Paired Student’s t-test; n ≥ 4 biological replicates. **(G-G’)** Immunofluorescence staining of ref(2)P and ubiquitin in the central brain at 7 and 13 dpe (G; scale bar = 50µm in main panels, 5µm in inset indicated by squared in the main panels) and puncta quantification (G’; data are shown as mean ± SEM. Unpaired Welch’s t-test; n = 3 biological replicates). **(H-H’)** GFP-mCherry-Atg8a reporter in the central brain at 13 dpe (H; scalebar 25µm). **(H’)** Quantification of mean mCherry and GFP fluorescence intensity (mean gray value; a.u.) and the GFP/mCherry mean gray value ratio (H’). Data are shown as mean ± SEM. Unpaired Welch’s t-test; n = 4 biological replicates. **(I-I’)** CathD-mCherry expression in the central brain at 13 dpe (I; scalebar 25µm). and quantification of mCherry-positive area relative to brain area in (I’; unpaired Welch’s t-test; n ≥ 5 biological replicates). **(J)** qPCR on head extracts of adult files carrying the genomic *CathD-mCherry* insertion and expressing 3R or 36R G4C2 repeats at 13 dpe. Gene expression is shown as mean fold change (2^−ΔΔCt^) relative to control. Data are shown as mean ± SEM. Paired Student’s t-tests performed on ΔCt values; n = 3 biological replicates. All experiments were performed using the *D42-GAL4* driver.

At the presymptomatic stage, qPCR analysis revealed coordinated transcriptional downregulation across multiple TFEB/Mitf target genes in 36R head extracts, when compared to 3R head extracts **(Figure 1B-E)**. Transcript levels of several lysosomal and vacuolar ATPase genes, including *emp, CathD*, *VhaAC45*, *Vha55*, *Vha100*-2, and *VhaSFD*, were significantly reduced by approximately 20-40% relative to 3R controls across biological replicates **(Figure 1D-E)**. Transcript levels of *Mitf* itself were also consistently decreased **(Figure 1B)**, suggesting early impairment of lysosomal transcriptional regulation. In contrast, early autophagy genes showed greater variability: *Atg8a* displayed a significant reduction, whereas *Atg9* and *ref(2)P* (encoding the autophagy adapter p62) exhibited variability across replicates **(Figure 1C)**.

Consistent with these observations, Western blot analysis of adult head extracts revealed a progressive accumulation of ref(2)P/p62, indicative of impaired autophagic cargo degradation in 36R animals, when compared to 3R **(Figure 1F-F’)**. In contrast, Atg8a protein levels, including both the non-lipidated (Atg8a-I) and lipidated (Atg8a-II) forms, as well as the Atg8a-II/Atg8a-I ratio, did not show statistically significant differences between genotypes at either age **(Figure 1F-F’)**.

To further assess cargo accumulation in the CB, we performed immunofluorescence staining to detect ref(2)P and ubiquitin, which marks cargoes recognized by ref(2)P. Already at the presymptomatic stage, 36R-expressing flies, but not 3R-expressing flies, exhibited the presence of colocalizing ref(2)P- and ubiquitin-positive puncta throughout the neuropil, which became more pronounced at the symptomatic stage **(Figure 1G-G’)**. These data indicate an early and progressive accumulation of autophagic and ubiquitinated cargoes.

We next examined the autophagic flux in G4C2-expressing neurons of the CB. To this end, we expressed a tandem *GFP-mCherry-Atg8a* reporter under the control of *D42-GAL4*, which exploits the differential sensitivity of GFP and mCherry to acidic environments to distinguish compartments **(Nezis et al., 2010).** At the symptomatic stage, fluorescence intensity analysis revealed increased mean GFP fluorescence in 36R-expressing CBs, while mean mCherry fluorescence was unchanged (**Figure 1H).** Consequently, the GFP/mCherry fluorescence-intensity ratio was significantly increased in 36R-, relative to 3R-expressing CBs (**Figure 1Hʹ**).

To assess the status of the lysosomal compartment in the CB of 36R-expressing flies, we examined animals also expressing *CathD-mCherry,* an endogenously expressed, tagged lysosomal-resident protease **(Takáts et al., 2013)**. Compared with age-matched 3R controls, 36R-expressing brains displayed a reproducible increase in the mCherry-positive puncta, with a trend already evident at the presymptomatic stage that reached statistical significance in symptomatic animals **(Figure 1I-I’)**. Notably, this occurred despite reduced *CathD* transcript levels in the heads of symptomatic flies **(Figure 1J)**.

Together, these data show that G4C2 repeat toxicity induces transcriptional suppression of lysosomal genes, progressive accumulation of ubiquitinated autophagy cargoes and alteration of the lysosomal compartment.

### Dual roles of the autophagy pathway in G4C2 toxicity

To assess the neurotoxicity associated to ALP defects, we analyzed the *Drosophila* compound eye, a well-established read out (**Bonini and Fortini, 2003)**. Consistent with previous studies, expression of 36R under control of *GMR-GAL4* resulted in the presence of necrotic patches in 8 dpe male flies **(Figure 2A) (Mizielinska et al., 2014)**. *Per se,* no downregulation using *GMR-GAL4* of any of the ALP genes tested resulted in formation of necrotic patches or lethality (not shown). However, downregulation of multiple lysosomal genes in flies expressing 36R, including *VhaSFD, Vha14*, *Vha100* and *Vha44,* enhanced necrotic patch formation and, in the extreme case of *CathD,* caused lethality **(Figure 2B**, quantified in **D).** These findings indicate that lysosomal degradation is protective against neurotoxicity and, considering the lysosomal disfunction observed in 36R animals, suggest that residual lysosomal is limited in 36R animals.

**Figure 2.**
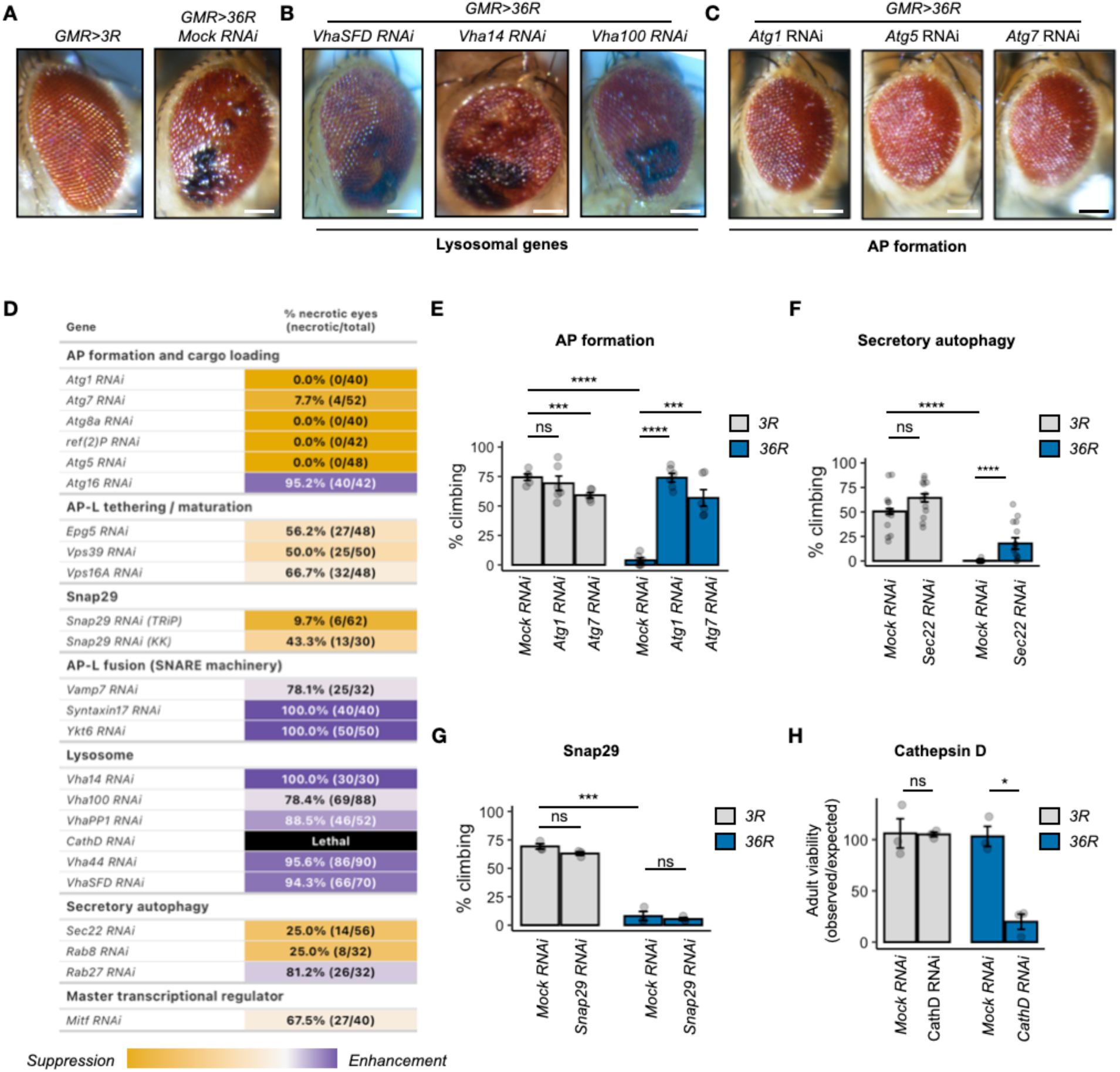
Genetic interaction analysis of ALP genes in G4C2-expressing flies. **(A-C)** Representative images of adult male eyes of *GMR>3R* animals or *GMR>36R* animals expressing a non targeting (mock) RNAi (A), or *GMR>36R* animals expressing RNAi to target the indicated lysosomal (B) or AP formation genes (C). Scalebars: 100µm. **(D)** Percentage and fraction of necrotic eyes for each listed downregulation in *GMR>36R* animals. Values represent pooled observations across independent crosses. The *GMR>36R* control resulting in 75% necrotic eyes represents the baseline. Genes are grouped by functional category. Shades of yellow indicate the extent of suppression, white indicates no modification relative to the baseline, shades of purple indicate enhancement and black lethality. **(E-G)** Percentage of normally climbing 13 dpe *D42>3R* or *D42>36R* flies expressing the indicated RNAi targeting AP formation (E; n ≥ 5 biological replicates) or secretory autophagy genes (F; n = 10 biological replicates) or *Snap29* (G; n = 3 biological replicates). Data are shown as mean ± SEM. Unpaired Welch’s t-test. **(H)** Percentage of survival to adulthood (number of observed animals/expected animals) of *D42>3R* or *D42>36R* flies expressing the *CathD RNAi*. Unpaired Welch’s t-test; n = 3 biological replicates.

Strikingly, reducing expression of genes encoding factors well known to act at early steps of autophagy, including *Atg1*, *Atg5*, *Atg7, Atg8a*, and *ref(2)P*, strongly suppressed necrotic patch formation, with the notable exception of *Atg16* **(Figure 2C**, quantified in **D)**. Thus, limiting the entry of cargoes into the autophagy pathway by preventing AP formation, alleviates neurotoxicity.

Downregulation of *Syntaxin17, Ykt6,* and *Vamp7,* which are involved in the AP-lysosome fusion enhanced necrotic patch formation, with only a mild effect observed for Vamp7 **(Figure 2D).** Thus, while AP biogenesis appears to be neurotoxic, AP fusion to lysosomes is neuroprotective.

Surprisingly, downregulation using two independent RNAi lines of *Snap29,* encoding the cytosolic SNARE also found to promote AP-lysosome fusion **(Mastrodonato et al., 2019; Morelli et al., 2014; Schiller et al., 2016)**, led to a suppression of G4C2 toxicity comparable to that observed upon depletion of early autophagy genes **(Figure 2D**). Partial suppression was also observed in flies heterozygous for a loss-of-function *Snap29* allele **(Figure S2A-A’)**. Because Snap29 is a multifunctional SNARE that acts in multiple membrane trafficking and non-trafficking pathways within the cell **(Morelli et al., 2014; Mastrodonato et al., 2018; Morelli et al., 2021; Smeele & Vaccari, 2023)**, we further dissected the contribution of Snap29-associated processes in neurotoxicity. To this end, we downregulated multiple known Snap29 interactors across distinct cellular pathways and measured 36R-induced necrotic patch formation. Also in this case, downregulation in the absence of 36R expression did not result in necrotic patch formation. However, downregulation of genes involved with Snap29 in ER/Golgi trafficking, endocytosis and non-trafficking functions predominantly enhanced 36R-associated degeneration phenotypes or caused lethality **(Figure S2B)**. Conversely, downregulation of genes implicated with Snap29 in the fusion of autophagy-related vesicles with the plasma membrane, including *Sec22* and *Rab8*, strongly suppressed degeneration **(Figure 2D)**.

To determine whether the suppression uncovered above extend to amelioration of motor impairment, we assessed the locomotor performance using the *D42-GAL4* driver. While downregulation of *Atg7* in 3R-expressing animals impairs locomotor function, recapitulating previous evidence **(Juhasz et al., 2007)**, downregulation of *Atg7* or *Atg1* in 36R-expressing animals prevented the climbing deficit observed in 36R-expressing flies at 13 dpe **(Figure 2E)**. A partial but highly significant rescue was also observed upon downregulation of *Sec22* **(Figure 2F)**. In contrast, *Snap29* downregulation did not rescue climbing behavior **(Figure 2G)**, suggesting that its protective effect in the eye may be context-dependent. As observed with *GMR-GAL4,* downregulation of *CathD* with *D42-GAL4* in the 36R animals significantly reduced adult viability **(Figure 2H)**, confirming the protective role of lysosomal degradation.

These genetic interaction data, when combined, reveal that while the activity of genes involved in lysosomal clearance is neuroprotective, that of genes that control AP formation and secretory autophagy contribute to neurotoxicity.

### Inhibition of autophagy and lysosomal activity differentially affect extracellular release of polyGA and autophagy-associated material

Prompted by our *in vivo* data, we next investigated how modulation of autophagy or lysosomal function affect trafficking of DPRs and autophagy-associated cargo *in vitro*. To isolate the effects of DPRs independent of G4C2 repeat RNA and RAN translation, we first used a codon-optimized, N-terminal FLAG-tagged polyGA construct in motor neuron-like NSC-34 cells. FLAG-polyGA is an aggregation-prone DPR that has previously been shown to impair lysosomal function **(Ferrari et al., 2026)**. Upon transfection, FLAG-polyGA was readily detected in both cell lysates and matched conditioned media (CM), demonstrating that polyGA is released into the extracellular environment under basal conditions **(Figure S3A).** In addition, we found FLAG-polyGA aggregates in the CM **(Figure S3B)**.

FLAG-polyGA-transfected cells were treated with either 3-methyladenine (3MA) to inhibit AP formation or NH₄Cl to impair lysosomal acidification and degradation. In line with previous studies (**Cristofani et al., 2018)**, 3MA treatment consistently increased the level of intracellular FLAG-polyGA, p62 and LC3-I **(Figure 3A)**. However, it did not change FLAG-polyGA, p62 or LC3 levels in the CM **(Figure 3B)**. In contrast, impairment of lysosomal degradation with NH₄Cl selectively increased the levels of the polyGA secreted in the CM, without a comparable increase in intracellular polyGA **(Figure 3A-B)**. In both control- and FLAG-polyGA-transfected cells, NH₄Cl also markedly increased the levels of p62 and LC3-I and LC3-II present in the CM, relative to untreated cells **(Figure 3B)**. GAPDH was undetectable in the CM **(Figure 3B)**, arguing against substantial contamination by bulk cytosolic release.

**Figure 3.**
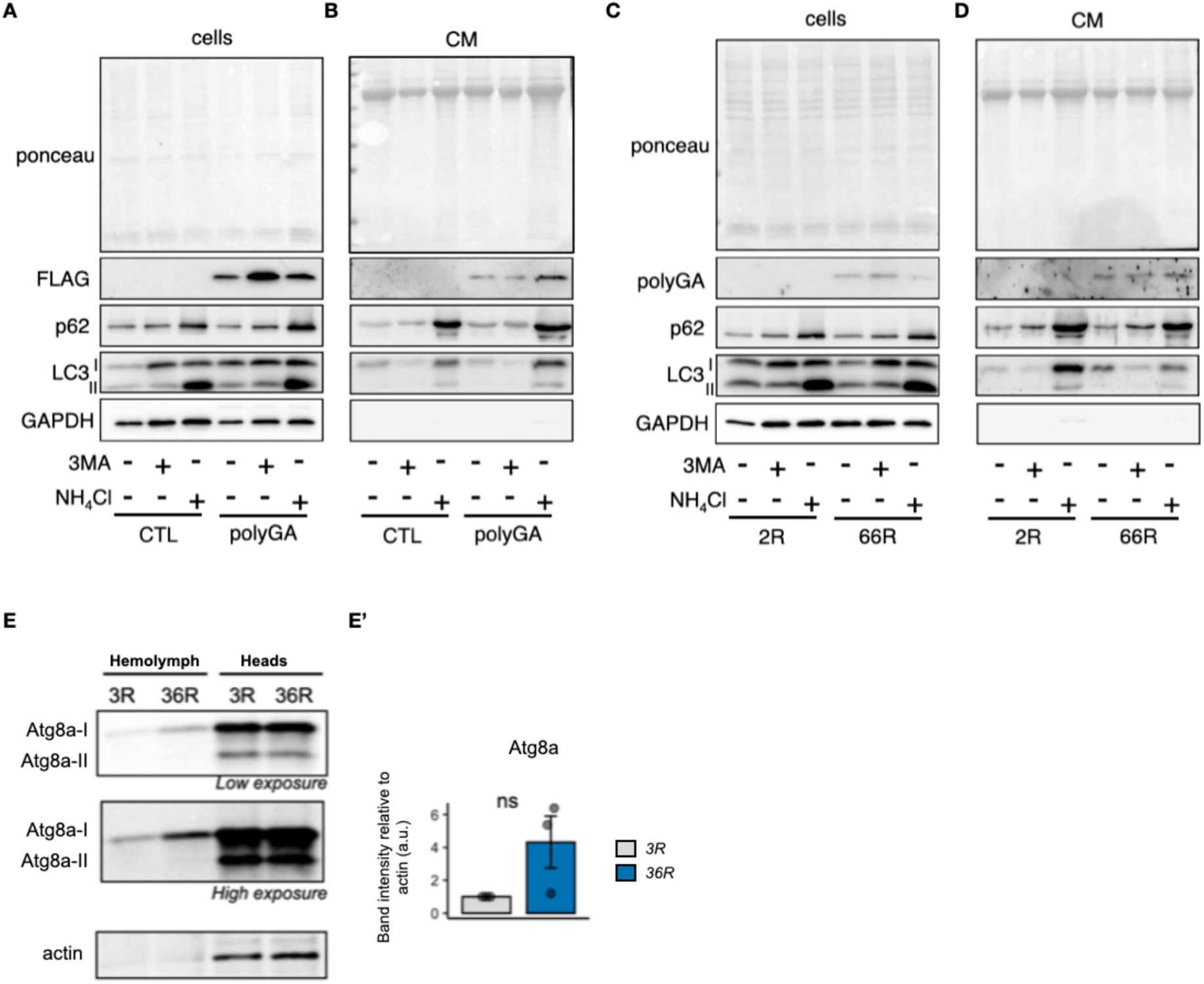
Lysosomal impairment causes extracellular release of DPRs and autophagy-associated proteins. **(A-B)** Immunoblotting of FLAG-polyGA, p62, LC3 and GAPDH in matched cell lysates (A; cells) and conditioned medium (B; CM) from NSC-34 cells transfected with pCDNA3 (CTL) or FLAG-polyGA and treated with 3MA, NH₄Cl or left untreated, as indicated. Ponceau staining is shown as a loading control for conditioned medium samples. **(C-D)** Immunoblotting of polyGA, p62, LC3 and GAPDH in matched cell lysates and conditioned medium from NSC-34 cells transfected with 2R or 66R G4C2 constructs and treated with 3MA, NH₄Cl or left untreated, as indicated. Ponceau staining is shown as a loading control for conditioned medium samples. **(E-E’)** Immunoblotting of Atg8a and actin in hemolymph and head extracts from 7dpe adult flies expressing 3R or 36R repeats under the *D42-GAL4* driver. Low and high exposure images are shown. Actin was readily detected in cell lysates but was essentially absent from hemolymph, arguing against nonspecific leakage as the primary source of extracellular Atg8a. Quantification of hemolymph Atg8a-I band intensity relative to actin (E’). Data are shown as mean ± SEM. Unpaired Welch’s t-test; n = 3 biological replicates.

We next asked whether these observations extend to a second type of disease-relevant G4C2 repeat. NSC-34 cells were transfected with either a non-pathogenic 2-repeat (2R) or pathogenic 66-repeat (66R) G4C2 construct, which undergoes RAN translation to generate the sense-derived DPRs polyGA, polyGP and polyGR **(Ash et al., 2013).** Detecting polyGA as a proxy for the DPRs produced by 66R, we first confirmed DPR presence and assessed lysosomal status. Compared with 2R-transfected cells, 66R-transfected cells exhibited increased GFP-GAL3 and LAMP1 expression 48 hours after transfection, consistent with lysosomal stress and the accumulation of lysosomes observed in the cell bodies of 36R-expressing flies **(Figure S3C).**

Then, we examined DPR levels in matched cell lysates and CM fractions. PolyGA was detected in both fractions from 66R-transfected cells but was absent from 2R-transfected cells. Similar to the FLAG-polyGA model, 3MA increased intracellular polyGA without increasing extracellular polyGA, whereas NH₄Cl selectively increased extracellular polyGA **(Figure 3C-D).** Thus, lysosomal impairment promoted extracellular polyGA release in both the codon-optimized and repeat-derived models. As in the FLAG-polyGA experiments, intracellular responses to both drugs were consistent with effective pathway inhibition: 3MA increased LC3-I and p62, whereas NH₄Cl increased LC3-II, LC3-I and p62 **(Figure 3C).** Analysis of the CM showed the same overall pattern observed in the FLAG-polyGA model **(Figure 3D)**.

We finally asked whether autophagy-associated cargo could also be detected extracellularly *in vivo*. To address this, we analyzed hemolymph from presymptomatic flies expressing 36R in neurons under the *D42-GAL4* driver. Low levels of non-lipidated Atg8a-I were found secreted in both 3R controls and 36R. 36R showed a trend towards increased Atg8a-I levels in hemolymph compared with 3R controls, although this difference did not reach statistical significance because of substantial variability between samples **(Figure 3E)**. Actin was low in hemolymph relative to head extracts, arguing against substantial contamination by intracellular material **(Figure 3E)**. As an additional control, GFP expressed under *D42-GAL4* was readily detected in head extracts but not in hemolymph **(Figure S3D)**, indicating that soluble neuronal proteins are not broadly released into the circulation. Since *D42-GAL4* is driven by a regulatory region associated with the *Toll-6* gene, we also examined *Toll-6* expression to assess whether the *D42* expression domain could include haemocytes. Analysis of Fly Cell Atlas data further showed little or no *Toll-6* expression in hemocytes, arguing against D42-driven expression in hemocytes as the source of the hemolymph Atg8a-I signal **(Figure S3E)**.

Together, these findings indicate that reduced lysosomal degradation, but not reduced autophagy induction is associated with increased extracellular release of DPR and autophagy-associated proteins.

### Poly-GR spreads to non-expressing glia and neurons in vivo

Since lysosomal dysfunction promoted extracellular release of DPRs and autophagy-associated proteins, we next asked whether this was accompanied by intercellular propagation of a toxic DPR *in vivo*. Consistent with our *in vitro* data, previous work showed that the polyGA spreads beyond neurons in which it is produced in the adult fly brain, when expressed in olfactory sensory neurons (OSNs) with the *Or83b-GAL4* driver **(Morón-Oset et al., 2019)**. To assess whether the more toxic polyGR is also released *in vivo*, we expressed a 44-repeat (44R) G4C2 RAN translated transgene containing polyGR in frame with GFP (GR-GFP).

We confirmed that 44R expression induces Mitf-dependent pathogenesis in salivary gland, which allow visualization of changes in the endogenous Mitf nuclear levels caused by modulation of regulators, such as 14-3-3ζ and RagA/B (**Figure S4A-A’**). Expression of 44R reduced nuclear endogenous Mitf levels (**Figure S4B-B’**), consistent with previous work (**Cunningham et al., 2020)**.

We next investigated GR-GFP localization in the CB, when expression is limited to OSN. When GFP is expressed in OSN in control animals, it is found confined to the expressing tissue **(Figure 4A)**. In sheer contrast, flies expressing 44R in OSN resulted in massive presence of GR-GFP signal concentrated in very large puncta throughout the CB, consistent with polyGR secretion from producing OSN, uptake and concentration in wide set of recipient cells **(Figure 4B**, quantified in **B’)**.

**Figure 4.**
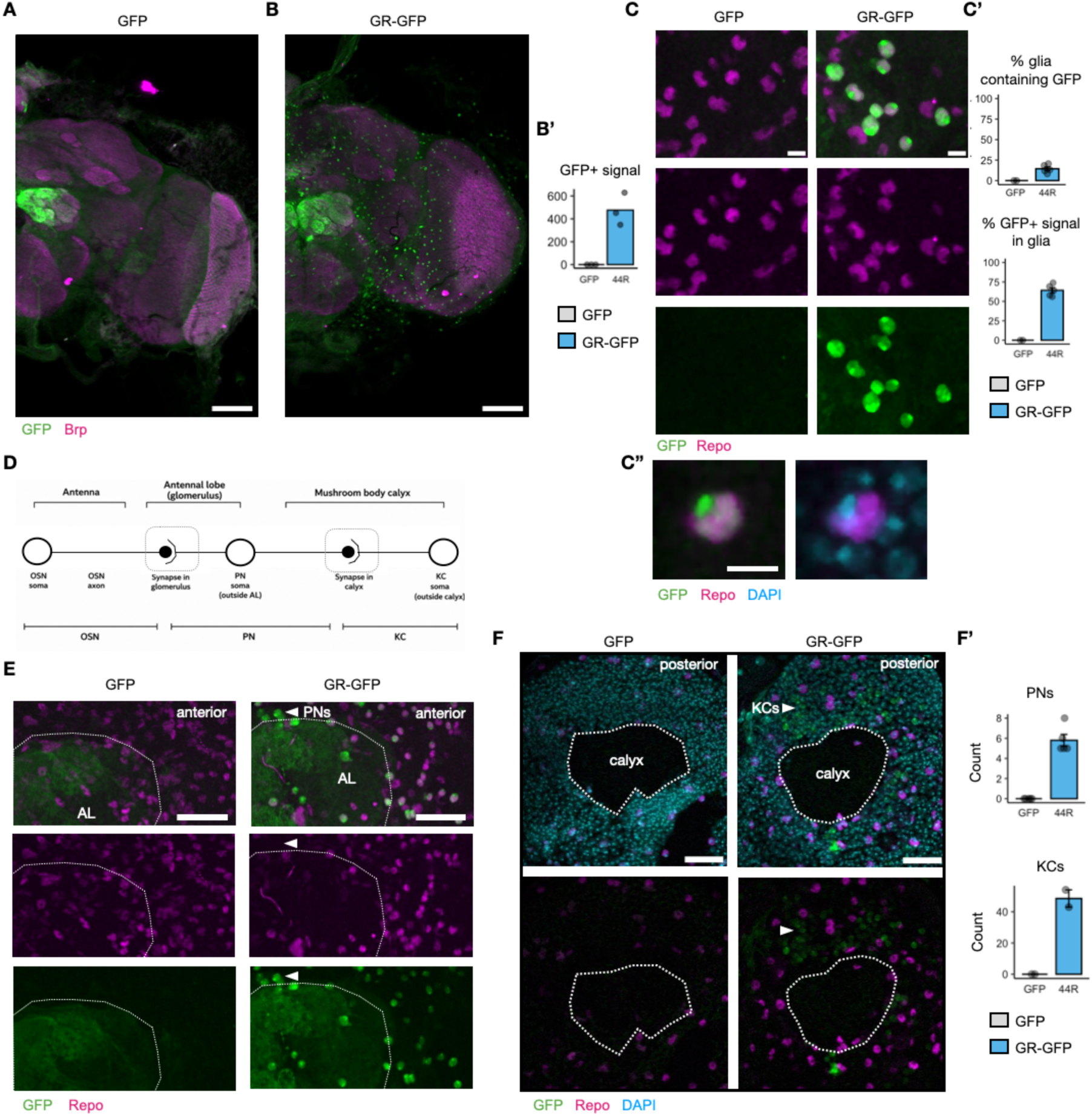
Poly-GR accumulates in non producing cells when expressed in OSNs. **(A-B’)** Representative images of adult fly brains expressing GFP or GR-GFP in OSNs under control of the *Or83b-GAL4* driver, immunostained as indicated (A-B; scale bars: 50µm). Count of GFP-positive puncta per brain. Data are presented as mean ± SEM (B’). **(C-C”)** Representative images of adult fly brains expressing GFP or GR-GFP in OSNs under control of the *Or83b-GAL4* driver, immunostained as indicated (C, C”; scale bars: 10µm). Quantification of GFP signal in glia. Data are presented as mean ± SEM (C’). **(D)** Schematics of the olfactory circuit indicating OSNs, connecting projection neurons (PNs), and connecting Kenyon cells (KCs). **(E-F’)** Representative images of anterior brain regions showing the AL in flies expressing GFP or GR-GFP with *Or83b-GAL4*), stained as indicated (E-F; scale bars = 20µm). Quantification of GFP-positive signal associated with PNs and KCs. Data are presented as mean ± SEM (F’).

We therefore sought to identify the recipient cell types as well as the subcellular site of accumulation. We first examined glial cells by co-staining with the glial nuclear marker Repo. While control flies expressing GFP did not show any signal in glial cells, a substantial proportion of the GFP signal produced by 44R was found within Repo-positive glial cells **(Figure 4C**, quantified in **C’**). Within these cells, the GFP signal was distributed throughout the nucleus with the strongest signal frequently observed in DAPI-dense and Repo-negative nuclear regions, suggestive of accumulation in heterochromatin **(Figure 4C”)**. Quantification across whole CB mounts showed that approximately 15% of glial nuclei contained GFP signal. When the analysis was restricted to the superficial ∼15 μm of the brain, a region enriched for cortex and surface-associated glia, this increased to approximately 36% of glial nuclei **(Figure S4C)**. A similar distribution was observed in the optic lobe, where the 44R-associated GFP signal accumulated predominantly in outer lamina-associated glia, whereas more medially located glia contained little or no detectable signal **(Figure S4D).**

Not all the GFP-positive signal in the CB of 44R animals was associated with glia. Around 40% was Repo-negative (**Figure 4C’**). We therefore asked whether signal could be detected in neurons connected to the OSNs. OSNs project to the antennal lobe, where they synapse onto projection neurons (PNs). PNs then relay olfactory information to higher-order brain centers, including the mushroom body calyx, where they synapse onto Kenyon cells (KCs) **(Figure 4D)**. In anterior 44R brain sections, the GFP-positive, Repo-negative signal was particularly evident in nuclei of neurons positioned around the antennal lobe, including in regions known to contain PN cell bodies **(Figure 4E)**. In posterior brain sections, similar GFP-positive signal was evident in nuclei positioned around the mushroom body calyx, including regions known to be enriched in KC soma **(Figure 4F**, quantified in **F’)**. No GFP signal outside the expression domain was detected in GFP-expressing control flies **(Figure 4E-F)**.

Together, these findings demonstrate that polyGR is secreted from producing neurons and accumulates in the nuclei of both glial cells and neurons within the olfactory circuit.

### VPA partly rescues G4C2-induced deficits in a stage-dependent manner

Considering the early reduction in Mitf activity and the lysosomal dysfunction inducing toxic DPR secretion, we next asked whether pharmacological activation of Mitf could ameliorate G4C2-induced phenotypes. Previous studies have shown that the FDA-approved VPA enhances TFEB-dependent transcription *in vitro* **(Kim et al., 2024)**. We therefore assessed whether chronic VPA treatment prevents the appearance of G4C2-induced phenotypes *in vivo*.

To examine effects on retinal degeneration, flies expressing 36R under the *GMR-GAL4* driver were raised on food containing 1 mM VPA throughout development. VPA supplementation markedly reduced the area of retinal necrosis in 36R-expressing flies, when compared with unsupplemented controls **(Figure 5A**, quantified in **Aʹ)**. We next asked whether this protective effect was dependent on *Mitf*. We found that the protective effect of VPA was significantly attenuated in flies depleted of *Mitf* **(Figure 5A**, quantified in **Aʹ)**. These findings indicate that Mitf contributes to the protective effect of VPA against retinal degeneration.

**Figure 5.**
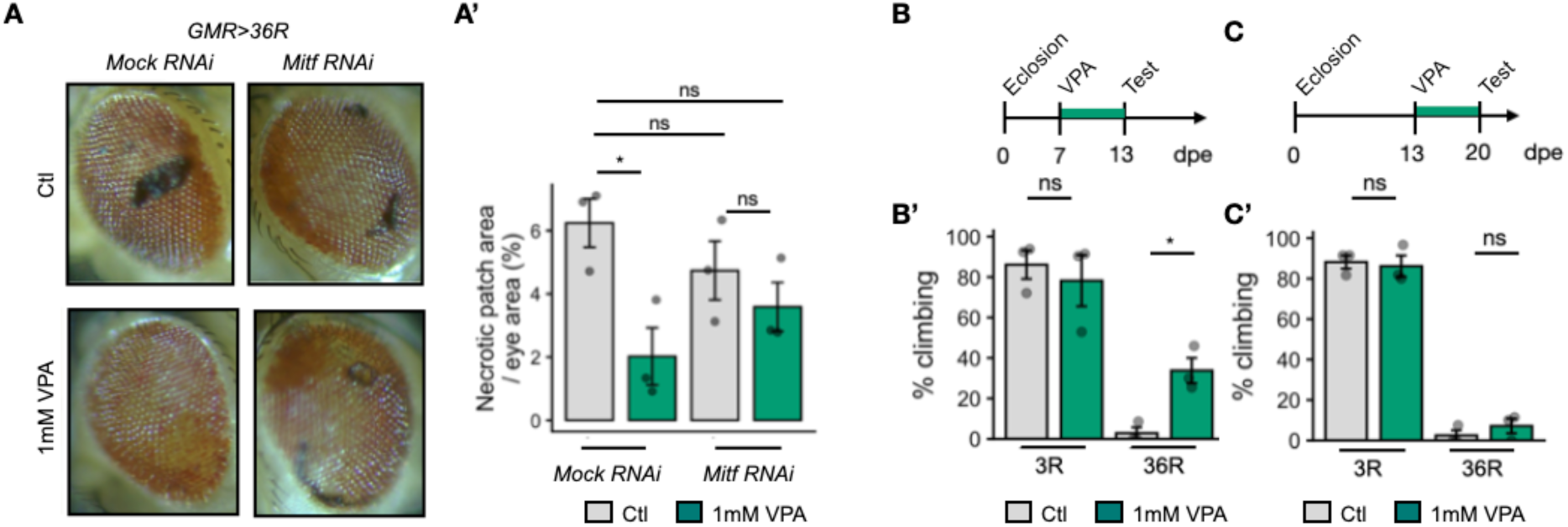
VPA prevents G4C2 repeat toxicity in *Drosophila*. **(A-A’)** Representative images of adult male eyes expressing 36R under *GMR-GAL4* and either mock RNAi or *Mitf* RNAi under standard food (Ctl) or food supplemented with 1 mM VPA (A). Quantification of necrotic patch area relative to total area in percentage. Each data point represents an average from >20 eyes (A’) Data are presented as mean ± SEM. Unpaired Welch’s t-tests. **(B-C)** Schematics of the experimental timelines for presymptomatic (B) and symptomatic (C) VPA supplementation, followed by assessment of climbing performance. **(B’-C’)** Climbing performance of flies expressing 3R or 36R with *D42-GAL4* following presymptomatic (B’) or symptomatic (C’) VPA supplementation. Statistical analysis was performed using unpaired Welch’s t-tests.

We next investigated whether VPA supplementation also improved the motor phenotypes of 36R flies and at which stage the supplementation is beneficial. When animals were supplemented with 1 mM VPA at the presymtomatic stage **(Figure 5B)**, VPA supplementation significantly increased the proportion of 36R flies reaching 3.5 cm within 10 s, compared with untreated controls **(Figure 5B’)**, indicating a partial improvement in locomotor function.

In contrast, when flies were supplemented with 1 mM VPA at the symptomatic stage **(Figure 5C**), although VPA-treated 36R flies appeared slightly more active than untreated controls, this improvement was not statistically significant **(Figure 5C’)**. Together, these findings show that VPA prevents the emergence of neurodegenerative phenotypes in the G4C2 model, but it is unable to revert phenotypes when locomotor deficits are apparent.

## Discussion

Multiple lines of evidence in mammalian and *Drosophila* models implicate dysfunction of the ALP in ALS/FTD, arising from both C9orf72 loss-of-function and/or gain-of-function toxicity associated to repeat RNA and DPRs **(Amick et al., 2016; Beckers et al., 2023; Boivin et al., 2020; Cunningham et al., 2020; Sullivan et al., 2016; Sung & Lloyd, 2023)**. Reduced nuclear localization of TFEB/Mitf **(Cunningham et al., 2020; Lin et al., 2024; Wang et al., 2023)** and downregulation of TFEB/Mitf target genes **(Xu et al., 2023)** have been proposed as cause of ALP dysfunction, yet when these defects arise relative to neuronal dysfunction and how is ALP impacted has remained unclear.

We show that selective transcriptional suppression of lysosomal genes occur at a presymptomatic stage in a *Drosophila* G4C2 model, preceding both behavioral decline and the accumulation of autophagic and ubiquitinated cargo. Together with previous reports of autolysosome accumulation in *Drosophila* **(Cunningham et al., 2020)**, these findings suggest that TFEB/Mitf impairment occurs before neurodegeneration and could be a target for intervention. Consistent with this, we find that pharmacological treatment using the clinically approved VPA partially ameliorates disease traits, if the intervention is applied before the onset of symptoms. This stage-dependent response is consistent with progressive impairment of Mitf nucleocytoplasmic trafficking and suggests that, at later disease stages, Mitf function may be too compromised to effectively prevent the effects of toxic DPR secretion. Although VPA has multiple molecular targets, our data support previous studies showing that it acts as a histone deacetylase (HDAC) inhibitor that enhances TFEB-dependent transcription **(Kim et al., 2024).** Thus, HDAC targeting could be more broadly explored as a preventive measure in familiar ALS.

Interestingly, VPA has previously been evaluated in two clinical trials in ALS but failed to demonstrate clear efficacy in broadly recruited patient cohorts (**Piepers et al., 2009; Boll M-C et al., 2025**). Given the marked molecular heterogeneity of ALS, however, there is increasing recognition that genetically stratified clinical trials may be required to identify therapeutic benefit within specific disease subtypes. VPA has also been investigated in combination with lithium, which activates TFEB through complementary signaling pathways, with encouraging results reported in a phase II ALS trial (**Boll M-C et al., 2025**). Our findings suggest that pharmacological enhancement of TFEB/Mitf activity may be particularly relevant for C9orf72-associated ALS/FTD, where impaired Mitf function appears to be an early pathogenic event rather than simply a downstream consequence of neurodegeneration. Whether combined activation of TFEB/Mitf more effectively restores lysosomal function and slows disease progression in C9orf72-associated ALS/FTD warrants further investigation.

In our hands, the genetic effects of TFEB/Mitf-dependent ALP disruption impact more lysosomal and V-ATPase genes than early autophagy genes, with Atg8a abundance and lipidation remaining largely unchanged in diseased animals. Together with our tissue analysis of ref(2)P, ubiquitin and lysosomal markers, these data indicate that AP formation is largely preserved while lysosomal degradative capacity is impaired. This distinctively pathologic ALP reorganization may reflect regulation by parallel stress-response pathways. In particular, NRF2/CncC induces p62/ref(2)P and subsets of ATG genes under oxidative stress (Pajares et al., 2016), a process relevant to C9orf72 pathology (**Au et al., 2024; Birger et al., 2019; Riemslagh et al., 2021)**. NRF2-dependent programs could partially compensate for TFEB/Mitf dysfunction, maintaining AP biogenesis even as lysosomal capacity becomes limiting.

The increased GFP/mCherry signal in autophagic flux experiments is consistent with accumulation of AP that cannot fuse with lysosomes. However, we have not tested lysosomal acidification/maturation, which could yield similar phenotypes. Consistent with this, we observe an expansion of the lysosomal comparments that could be due an accumulation of dysfunctional lysosomes. Our transcriptional data exclude that this expansion is due to increased lysosome biogenesis. Cargo accumulation further supports a defect in degradative efficiency rather than AP formation. Such lysosomal phenotypes are well-established hallmarks of lysosomal storage diseases (LSD), such as Gaucher disease and Niemann-Pick’s disease, whereby mutations in lysosomal enzymes lead to progressive accumulation of undegraded substrates within dysfunctional lysosomes **(Platt et al., 2018)**. Although C9orf72-associated ALS is not strictly an LSD, our dissection of similar lysosomal defects may be relevant for future therapeutic strategies. Lysosomal stress phenotypes have been described across C9orf72 models, including impaired lysosomal acidification and maturation in mammalian neurons **(Beckers et al., 2023; Boivin et al., 2020)** and accumulation of autolysosomes in *Drosophila* **(Cunningham et al., 2020; Xu et al., 2023)**. Together with these, our observations points at limiting lysosomal degradative capacity as a key vulnerability associated to early G4C2 toxicity.

The imbalance between preserved AP formation and impaired lysosomal degradation raises a key question: how are autophagy-associated cargoes handled when degradative capacity becomes limiting? We propose that a fraction of cargoes including DPRs is redirected toward autophagy-dependent secretion, which has been implicated in the extracellular release and intercellular propagation of aggregation-prone proteins across multiple neurodegenerative contexts **(Nilsson et al., 2013; Kang et al., 2019; Hutchings et al., 2024; Nakamura et al., 2024; Sawai et al., 2025)**. Lysosomal dysfunction is a known trigger of secretory autophagy **(Leidal et al., 2020; Solvik et al., 2022)** and the outcome of our genetic modifier screen is consistent with such a shift. Reduction of genes supporting lysosomal function or AP-lysosome fusion enhances G4C2-induced degeneration, indicating that degradative flux is protective. In contrast, inhibition of early AP formation suppresses toxicity, as does depletion of genes associated with a non-canonical, secretory branch of the autophagy pathway. Together, these opposing genetic effects indicate that, under conditions of impaired lysosomal function, autophagy is not simply defective but functionally rerouted in a manner that contributes to toxicity.

Our genetic findings are consistent with previous *Drosophila* studies showing that reduction of ref(2)P/p62 suppresses G4C2 toxicity whereas its overexpression enhances degeneration **(Cunningham et al., 2020)**. In contrast, Xu et al. reported that *Atg1* downregulation worsened survival in an inducible neuronal G4C2 model already displaying early locomotor impairment **(Xu et al., 2023)**, suggesting that the impact of autophagy modulation depends on disease stage. Early suppression of AP formation may be beneficial when lysosomal flux is limiting, but detrimental once pathology is advanced. These findings highlight the importance of temporal context when interpreting autophagy manipulations.

Consistent with rerouting of cargo trafficking under conditions of lysosomal dysfunction, we found that impairment of lysosomal degradation promoted extracellular accumulation of both DPRs and p62 in two independent cellular models. Importantly, inhibition of AP formation increased intracellular polyGA without increasing its extracellular accumulation, demonstrating that intracellular DPR burden alone is insufficient to drive extracellular release.

Extending *in vitro* findings *in vivo,* we show that the toxic polyGR spreads beyond its neuronal expression domain and accumulates in both glial cells and neurons. Within the olfactory system, polyGR-associated signal was detected in neurons downstream of the expressing populations, raising the possibility that secretion and subsequent uptake of toxic DPRs could contribute to this progressive dissemination of pathology. In addition, polyGR preferentially accumulated in surface-associated glia, which contribute scavenging and blood-brain barrier-like functions in the *Drosophila* brain. Interestingly, we show that neuronal G4C2 toxicity is associated with high levels of Atg8a in the hemolymph of presymptomatic flies. Atg8a and its orthologs have been shown to be secreted by autophagy-dependent secretion upon lysosome inhibition (**Xu et al., 2022)**. This evidence raise the possibility that secretory autophagy may promote the propagation of DPRs to glia via the brain-hemolymph interface. However, whether neuronal propagation, glial accumulation, or both arise through altered autophagy-associated trafficking remains to be determined.

Interestingly, transferred polyGR-GFP frequently accumulated within intensely DAPI-positive regions of recipient glial nuclei. Although the identity of these nuclear domains remains unresolved, arginine-rich DPRs have previously been shown to associate with heterochromatin and perturb chromatin organization **(Zhang et al., 2019)**, raising the possibility that secreted polyGR, when uptaken by glia, might disrupt specific nuclear compartments.

Our findings differ from those of **Morón-Oset et al. (2019),** which reported propagation of the relatively non-toxic DPR polyGA, but not polyGR, in the adult *Drosophila* brain. An important distinction is that Morón-Oset et al. examined individually expressed, codon-optimized DPRs, whereas our study uses a G4C2 construct that generates all sense DPRs in the context of the repeat RNA. This more closely reflects the disease context and may influence intercellular propagation. Interestingly, polyGA propagation occurred in the absence of overt neuronal loss **(Morón-Oset et al., 2019).** However, polyGA is largely non-toxic in *Drosophila*, and whether propagation of the highly toxic DPR polyGR likewise occurs independently of neurodegeneration remains to be determined.

Collectively, our data support a model in which early TFEB-dependent lysosomal dysfunction is a central feature of C9orf72 repeat toxicity. Rather than impairing induction of autophagy, lysosomal degradative capacity becomes limiting while AP formation is largely preserved, diverting APs containing toxic DPRs towards secretion. A compound with HDAC inhibitory activity, previously shown to increase of TFEB activity, lead to reduced DPR toxicity, providing a promising therapeutic strategy to possibly prevent the mistrafficking events associated with C9orf72 ALS/FTD.

## Materials and Methods

### Drosophila maintenance and husbandry

Flies were maintained on standard cornmeal-yeast-agar medium supplemented with antimicrobials at appropriate concentrations to prevent bacterial and fungal contamination. Where indicated, food was supplemented with valproic acid (VPA) at a final concentration of 1 mM, as previously described (Yi et al., 2013). Flies were kept under controlled conditions at 25 °C, unless otherwise specified.

The *D42-GAL4* (BL8816), *GMR-GAL4* (BL9146) and *O83b-GAL4* (BL23292) driver lines and UAS-G4C2 repeat expansion lines carrying 3, 36, or 44 G4C2 repeats (*UAS-G4C2 3R* BL58688, *UAS-G4C2 36R* BL58687, and *UAS-G4C2 44R* BL4727), were obtained from the Bloomington Drosophila Stock Center. Additional UAS-RNAi, UAS-overexpression, and reporter lines used in this study are listed in **Supplementary table 1**. UAS-RNAi lines were selected based on published literature showing functional validation (e.g. *Atg1* RNAi **(Zirin et al., 2013)**, *Atg7* RNAi **(Andresen et al, 2025)**, and *Mitf* RNAi **(Bouche et al., 2016)**)

For experiments requiring multiple genetic manipulations, recombinant or double-balanced stocks carrying both the *GAL4* driver and *UAS-G4C2* transgene were maintained over a *CyO-GAL80* balancer to suppress GAL4/UAS expression during stock maintenance. These stocks were subsequently crossed to the appropriate UAS-RNAi, UAS-overexpression, or reporter lines to generate experimental progeny.

After eclosion, the adult progeny was transferred to 29 °C to enhance GAL4/UAS-driven transgene expression. Flies were maintained on fresh food flipped every 2-3 days. All experiments were performed on age-matched adult flies reared under non-crowded conditions. Unless otherwise specified, equal numbers of male and female flies were used for all analyses.

### Drosophila phenotypic assays

To score eye degeneration, male flies were analyzed at 8 dpe. Eyes were scored blind for the presence of necrotic patches by visual inspection. The total number of eyes with necrotic patches was quantified across three independent crosses and pooled for analysis.

For more sensitive analysis, the area occupied by necrotic patches was quantified as a percentage of the total eye area.

To evaluate climbing performance, flies were housed together for 2-3 days following eclosion to allow mating. Females were separated under brief CO₂ anesthesia and transferred to fresh food at a density of 6-13 flies per vial, with no further CO₂ exposure. Climbing assays were performed at 7, 13 or 20 dpe, as indicated. Each vial was subjected to five consecutive trials (technical replicates) with a 5 min rest interval between trials. Unless otherwise stated, climbing performance was assessed as the proportion of flies climbing above 7 cm within 10 s. Values were averaged across technical replicates. The indicated number of biological replicates were generated from independent crosses.

To assess viability, crosses were set using 4-6 virgin females and 3 males per vial and allowed to mate for 3 days. Vials were then flipped every 2 days for a week, generating 3-4 technical replicates per cross. The number of adult progeny of each genotype was recorded daily until all pupae had eclosed or for a maximum of 5 days. Viability was calculated as the ratio of observed to expected progeny based on Mendelian inheritance. Values were averaged across technical replicates. The indicated number biological replicates were generated from independent crosses.

### Cell culture, transfections and pharmacological treatment

The mouse motor neuron-like hybrid cell line NSC34 was cultured in high-glucose DMEM (EuroClone, ECB7501L) supplemented with 5% fetal bovine serum (Sigma-Aldrich, F7524), 1 mM L-glutamine (EuroClone, ECB3004D), and penicillin-streptomycin (EuroClone, ECB3001D) at 37 °C in a humidified atmosphere containing 5% CO₂. Cells were plated at a density of 80,000 cells/mL and transfected with 1 μg plasmid DNA using Lipofectamine™ 3000 (Thermo Fisher Scientific, L3000015) according to the manufacturer’s instructions.

Cells were transfected with *pAG3-CAG-(G4C2)₂ (2R)* or *pAG3-CAG-(G4C2)₆₆ (66R)*, a FLAG-tagged codon-optimized polyGA expression plasmid (100 repeats) kindly provided by Professor Daisuke Ito (Keio University School of Medicine, Tokyo, Japan) (Yamakawa et al., 2015) or the empty pcDNA3 vector as a transfection control. To study G4C2 impact on lysosome cells were co-trasfected with *EGFP-LGALS3* plasmid was kindly provided by Prof. M. A. Jäättelä (Danish Cancer Society Research Center, Copenhagen, Denmark).

Where indicated, cells were treated with 3-methyladenine (3MA; 10 mM) for 24 h or ammonium chloride (NH₄Cl; 4 mM) for 16 h prior to collection.

### Immunofluorescence staining, imaging and analysis

Adult fly brains and larval salivary glands were dissected in ice-cold PBS and fixed in 4% formaldehyde for 20 minutes at room temperature. Fixed tissues were washed three times for 5 minutes in PBS and permeabilized in PBS containing 0.3% Triton X-100 (PBST) for 1 hour at room temperature. Tissues were then blocked for 1 hour in 5% bovine serum albumin diluted in PBST.

Brains and larval salivary glands were incubated with primary antibodies diluted in blocking solution for 48 hours or 24 hours, respectively, at 4 °C on a rotator. Primary antibodies used were anti-ref(2)P (Abcam, ab178440; 1:500), anti-FK2 (Enzo Life Sciences, BML-PW8810; 1:150), anti-Brp (Developmental Hybridoma Bank, nc82, 1:40), anti-mCherry (1:500), anti-GFP (Abcam, ab13970, 1:400), and anti-Mitf (custom-made, **Tognon et al., 2016**, 1:500). Following primary antibody incubation, tissues were washed three times for 10 minutes in PBST and incubated with Alexa Fluor-conjugated secondary antibodies (1:500). Secondary antibody incubation was performed for 48 hours at 4 °C for adult brains and for 2 hours at room temperature for larval salivary glands. Unless otherwise indicated, fluorescently tagged reporters were directly visualized without staining.

Confocal images were acquired using a Zeiss LSM510 confocal microscope. Z-stack images were collected using identical acquisition settings for all genotypes within each experiment and processed as maximum-intensity projections for image analysis. Confocal stacks for figure 4 were acquired using Zeiss LSM800 confocal microscope, using a Plan-Apochromat 40x/1.3 oil-immersion objective.

Regions of interest were generated using Gaussian blur followed by automated intensity thresholding using the Otsu algorithm. Thresholds were applied identically across all genotypes within each experiment. All quantitative measurements were performed on unprocessed maximum-intensity projections.

### Protein extraction and immunoblotting

For total protein extracts, ten heads (5 female and 5 male) were homogenized in 100 µL of 1× Laemmli buffer (BIO-RAD, #1610747). Homogenates were boiled at 98 °C for 5 minutes and centrifuged at 14,000 × *g* for 10 minutes. Supernatants were collected and stored at -80 °C until use. For SDS-PAGE, 15 µL of each sample was loaded onto 4-20% Mini-PROTEAN TGX Precast Protein Gels (BIO-RAD, #4561094).

Hemolymph extraction was performed as previously described (**Damrau et al., 2017**), with minor modifications. Briefly, a hemolymph collection device was prepared by puncturing the bottom of a 500µL microcentrifuge tube with a 25G needle to create a single hole and placing this tube into a 1.5 mL microcentrifuge tube. The assembly was kept on ice at all times. Adult flies were handled individually and pierced once in the thorax using a 0.1 mm minutien pin. Flies were held by the wings to minimize additional tissue damage. Each punctured fly was immediately placed into the hemolymph collection device on ice. Samples were centrifuged at 5,000 × g for 10 minutes at 4 °C to collect hemolymph. Ice-cold RIPA buffer (Thermo Fisher Scientific, #89901) supplemented with 1× protease inhibitor cocktail (Merck, #04693132001) was added to the collected hemolymph, and samples were stored at -80 °C.

Protein concentrations were estimated using the Pierce BCA Protein Assay Kit (Thermo Fisher Scientific, A55864). For SDS-PAGE, 13ug of protein, were loaded onto 4-20% gradient gels. Following electrophoresis, proteins were transferred onto nitrocellulose membranes using the Trans-Blot Turbo Transfer System (BIO-RAD, 1704150EDU) according to the manufacturer’s instructions. Membranes were blocked for 1 hour at room temperature in 5% milk (Merck, #70166) diluted in TBS (BIO-RAD, #1706435) containing 0.1% Tween-20 (BIO-RAD, #1662404). After blocking, membranes were incubated overnight at 4 °C with primary antibodies against actin (Developmental Hybridoma Bank, JLA20, 1:10,000), ref(2)P (Abcam, AB178440, 1:500), or Atg8a (Abcam, AB109364, 1:1,000). The following day, membranes were washed three times in TBST and incubated with HRP-conjugated secondary antibodies (Cell Signaling Technology, 7076S and 7074S, 1:3,000) for 2 hours at room temperature. Where necessary, membranes were stripped and blocked in 5% milk before reprobing.

For NSC34 cell extracts, cells were collected 48 h after transfection. Conditioned medium was collected and centrifuged at 500 × g for 10 min and 2000 x g for 20 min to remove cellular debris. Extracellular proteins were concentrated by trichloroacetic acid (TCA) precipitation. Briefly, TCA was added to a final concentration of 20%, samples were incubated for 1h, centrifuged at 16000 × g for 10 min, washed twice with ice-cold acetone, air-dried, and resuspended in 1× Laemmli buffer.

Cells were washed once with PBS and collected by centrifugation at 100 × g for 5 min at 4 °C. Cell pellets were resuspended in PBS supplemented with 1× protease inhibitor cocktail (Sigma-Aldrich, P8340) and lysed by sonication. Protein concentrations were determined using the Quantum Protein Assay Kit (Cyanagen PRTD1,0500).

For SDS-PAGE, 20 μg of total protein or equivalent volumes of concentrated conditioned medium were loaded onto 12% polyacrylamide gels. Following electrophoresis, proteins were transferred onto nitrocellulose membranes (Amersham Protran, GE Healthcare) using a trans-Blot apparatus (Mini Trans-Blot Cell; Bio-Rad Laboratories) according to the manufacturer’s instructions. Membranes were blocked for 1 h at room temperature in 5% milk diluted in TBS containing 0.1% Tween-20 (TBST) and incubated overnight at 4 °C with primary antibodies against FLAG ( Sigma-Aldrich, F1804, 1: 1000), polyGA (Millipore; MABN889, 1:1000), p62 ( Sigma-Aldrich, P0067, 1:1000), LC3 ( Sigma-Aldrich, L8918, 1:1000), GAPDH ( Immunological Sciences, MAB-10578, 1:3000), LAMP1 ( Abcam, AB24170; 1:1,000), and GFP (<u>Immunological Sciences</u>, MAB-94345, 1:1000), as indicated. Membranes were washed three times in TBST and incubated with HRP-conjugated secondary antibodies anti-mouse IgG-HRP ( Jackson ImmunoResearch, 115–035-003, 1:5000) or anti-rabbit IgG-HRP (Jackson ImmunoResearch, 111-035-003, 1:5000) for 1 h at room temperature. Immunoreactive bands were visualised using enhanced chemiluminescence with kit reagents ECL Antares (Cyanagen, XLS142) or Westar hypernova (Cyanagen, XLS149). Where necessary, membranes were stripped and reprobed following blocking in 5% milk.

To detect insoluble polyGA species, filter trap assays were performed. Equal amounts of total protein (6 μg) were filtered through a 0.22 μm cellulose acetate membrane (Whatman GE Healthcare, GEH10404180) using a Bio-Dot SF Microfiltration Apparatus (Bio-Rad Laboratories, 1703938). Membranes were processed as described above for immunoblotting and incubated with either anti-FLAG or anti-polyGA primary antibodies, as indicated.

### RNA extraction and RT-qPCR

RNA was extracted from 20 adult fly heads (10 female and 10 male) using TRIzol™ Reagent (Thermo Fisher Scientific, #15596026) according to the manufacturer’s protocol, with minor modifications. Fly heads were homogenized in 500 µL of TRIzol using a pestle and incubated at room temperature for 5 minutes. Phase separation was achieved by adding 100 µL of chloroform, vortexing vigorously, and centrifuging at 11,000 × *g* for 15 minutes. The aqueous phase was carefully collected, and chloroform extraction was repeated once to improve purity. RNA was precipitated by adding an 0.5:1 isopropanol and incubating overnight at -80°C.

The following day, samples were centrifuged at 11,000 × *g* for 15 minutes and RNA pellets were washed with ice-cold 75% ethanol, air-dried briefly, and resuspended in RNase-free water. RNA concentration and purity were determined using the NanoDrop One 21 (Thermo Fisher Scientific). cDNA was synthesized from 800 ng of total RNA using the LunaScript RT SuperMix (New England Biolabs, M3010L) following the manufacturer’s instructions. Quantitative PCR was performed using LUNA Universal qPCR Master Mix (New England Biolabs, M3003E) using 12 ng of cDNA per reaction with gene-specific primers listed in **Supplementary table 2**.

### Statistical analysis

Data and statistical analyses were performed in R. Data are presented as mean ± SEM, unless otherwise stated. Comparisons between two independent groups were performed using two-sided unpaired Welch’s t-tests. Paired Student’s t-tests were used for paired experimental data, including RT-qPCR analyses performed on ΔCt values and the indicated immunoblot analyses. Comparisons among three or more groups were performed using one-way ANOVA followed by Tukey’s multiple-comparisons test, where indicated. The statistical test used for each experiment and the number of biological replicates are specified in the corresponding figure legend. Differences were considered statistically significant at P < 0.05. Significance is indicated as follows: ns, P ≥ 0.05; *, P < 0.05; **, P < 0.01; ***, P < 0.001; and ****, P < 0.0001.

## Author Contributions

Conceptualization, P.H.S. and T.V. Methodology, P.H.S., V.F., A.P. and T.V.; formal analysis, P.H.S., V.F., G.S., E.D., A.P. and T.V.. Resources, E.D., R.C., A.P. and T.V.. Funding acquisition, R.C. A.P. and T.V. Writing—original draft preparation P.H.S. and T. V. Review & editing, all authors.

## Abbreviations

ALP: autophagy-lysosome pathway
ALS: amyotrophic lateral sclerosis
AP: autophagosome
CB: central brain
CM: conditioned medium
C9orf72 (C9): chromosome 9 open reading frame 72
dpe: days post-eclosion
DPR: dipeptide repeat protein
FTD: frontotemporal dementia
G4C2: GGGCC
HDAC: histone deacetylase
KC: Kenyon cell
LSD: lysosomal storage disease
NRF2 (CncC): nuclear factor erythroid 2-related factor 2 (Drosophila Cap ‘n’ Collar C)
OSN: olfactory sensory neuron
PN: projection neuron
RAN: repeat-associated non-AUG (translation)
ref(2)P: refractory to sigma P
SNARE: soluble N-ethylmaleimide-sensitive factor attachment protein receptor
TFEB (Mitf): transcription factor EB (Drosophila microphthalmia-associated transcription factor)
V-ATPase: vacuolar-type H+-ATPase
VNC: ventral nerve cord
VPA: valproic acid.

## Supplementary Figures

**Figure S1.**
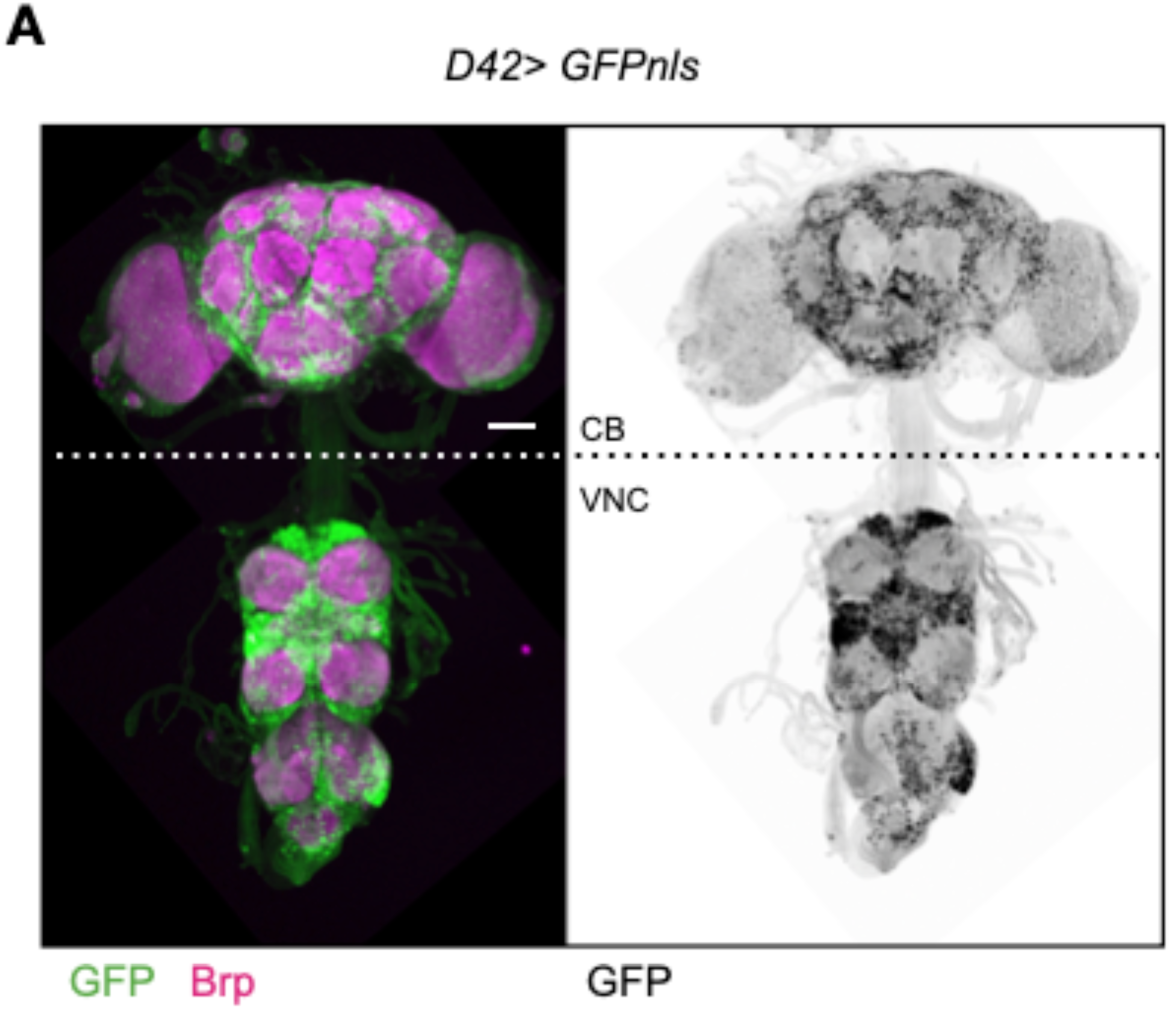
*D42-GAL4* expression in CB and VNC. **(A)** Representative images of the adult nervous system, comprising the central brain with optic lobes (CB) and the ventral nerve cord (VNC), showing nuclei of D42-expressing cells labeled with nuclear GFP (D42>GFP_nls_) in flies aged 7 dpe. Scalebar 25µm.

**Figure S2.**
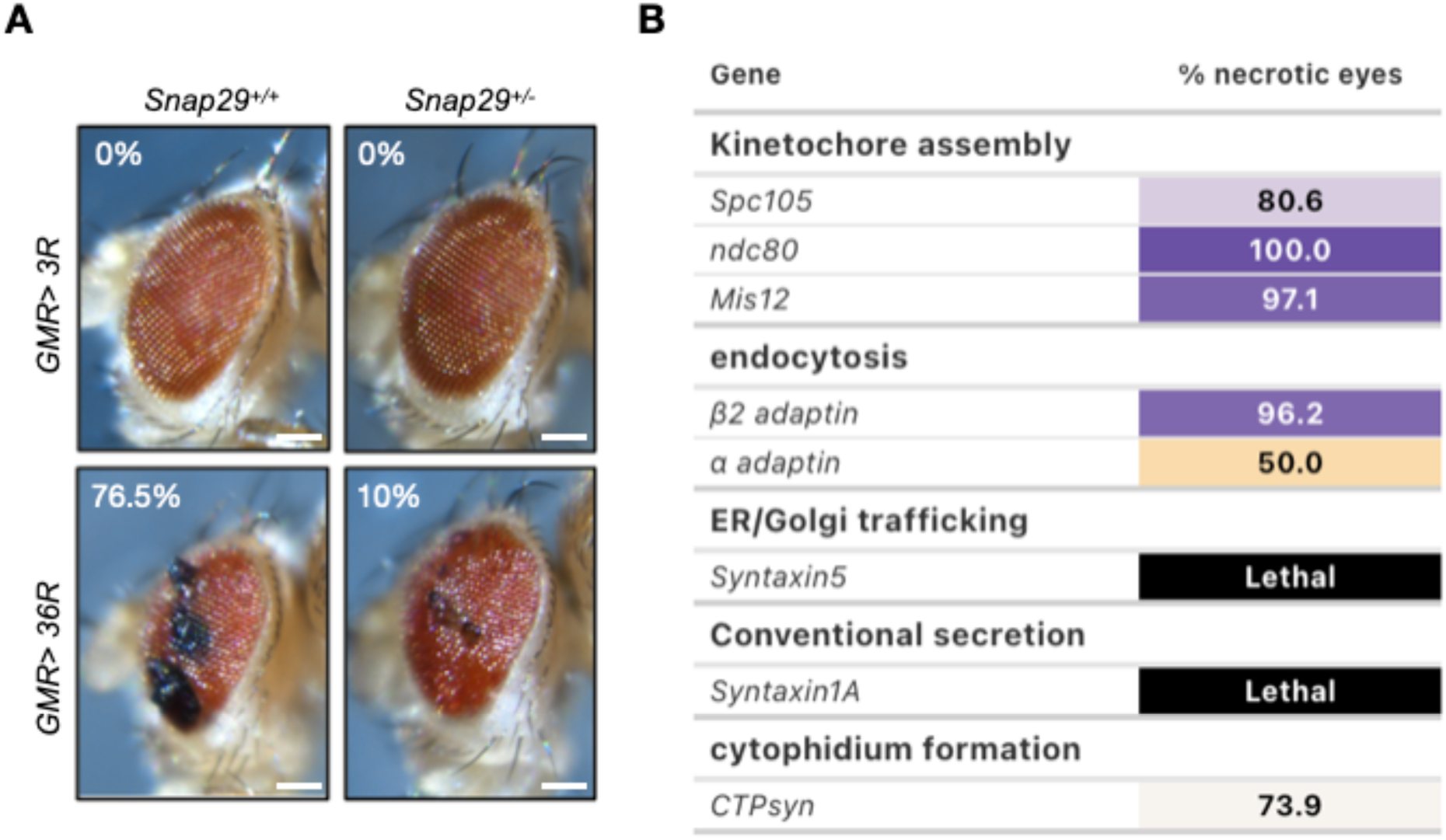
Genetic interaction analysis of *Snap29* and interacting genes. **(A)** Representative images of adult male eyes of *GMR>3R* animals or *GMR>36R* animals expressing in a Snap29^+/+^ or Snap29^+/-^ background, as indicated. Quantification is shown as percentage of eyes with necrotic patches (n>20). Scalebars: 100µm. **(B)** Percentage of necrotic eyes for each listed downregulation in *GMR>36R* animals. The *GMR>36R* control represents the baseline. The targeted *Snap29*-interacting genes are grouped by functional category. Shades of yellow indicate the extent of suppression, white indicates no modification relative to the baseline, shades of purple indicate enhancement and black lethality.

**Figure S3.**
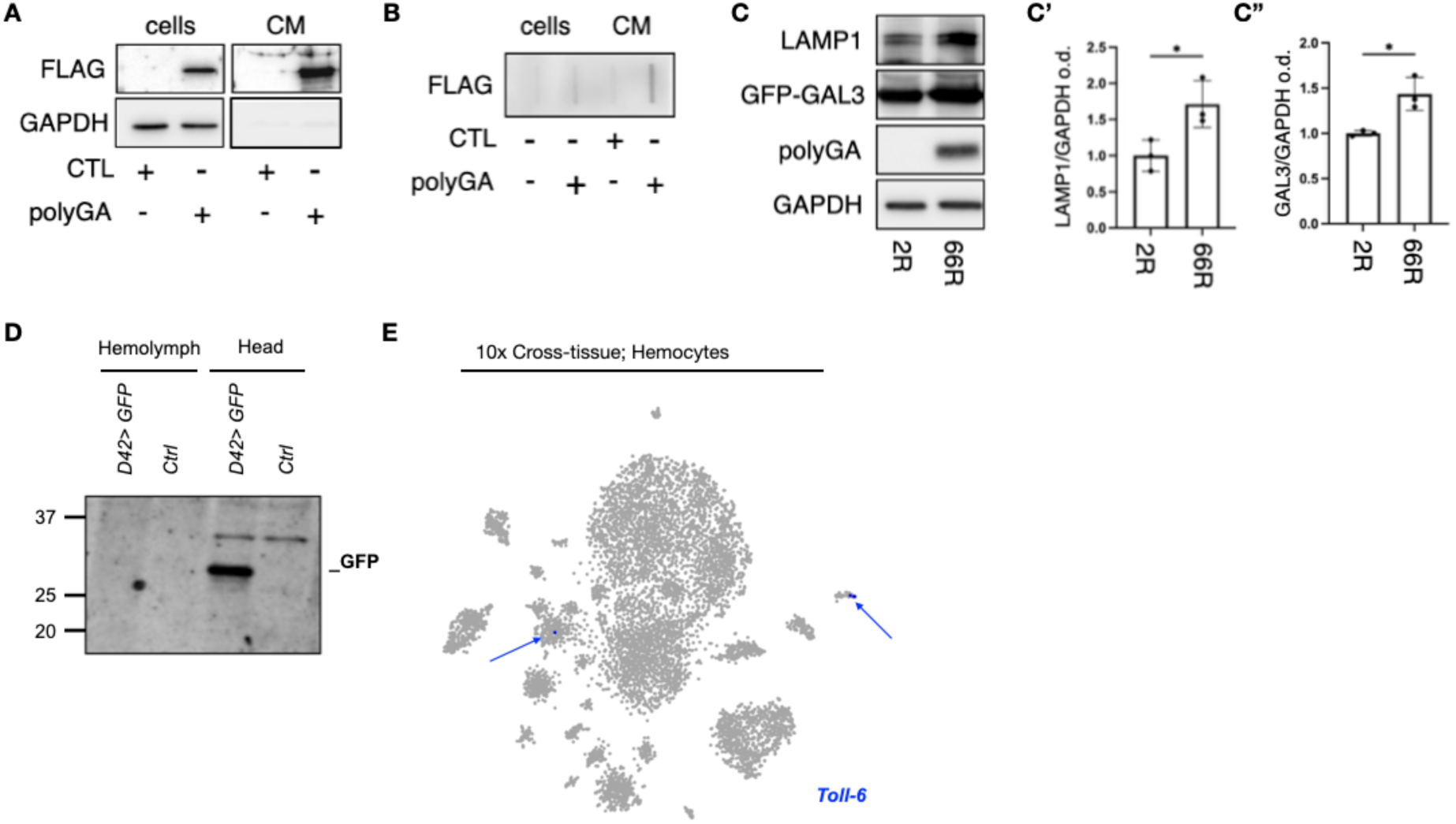
DPR and GFP expression controls. **(A)** Immunoblotting of FLAG-polyGA in cell lysates (cells) and conditioned medium (CM) from NSC-34 cells transiently transfected with pCDNA3 (CTL) or FLAG-polyGA, as indicated. GAPDH was used as a loading control for cell lysates. **(B)** Filter-trap assay detecting aggregated FLAG-polyGA in cell lysates and conditioned medium from NSC-34 cells transfected with pCDNA3 (CTL) or FLAG-polyGA. **(C-C”)** Immunoblotting of LAMP1, GFP-GAL3, polyGA and GAPDH in NSC-34 cells transfected with non-pathogenic (G₄C₂)₂ or pathogenic (G₄C₂)₆₆ repeat constructs (C). Quantified band intensity relative to actin in (C’) for LAMP1 and (C’’) for GAL3. Data are shown as mean ± SD. Unpaired Student’s t-test; n = 3 biological replicates. **(D)** Representative immunoblotting showing GFP expression in *D42>GFP* head lysates but not in hemolymph extracts. Molecular weight markers (kDa) are indicated. **(E)** UMAP projection of hemocytes from the FlyCellAtlas cross-tissue single-cell RNA sequencing dataset showing D42/*Toll-6* expression.

**Figure S4.**
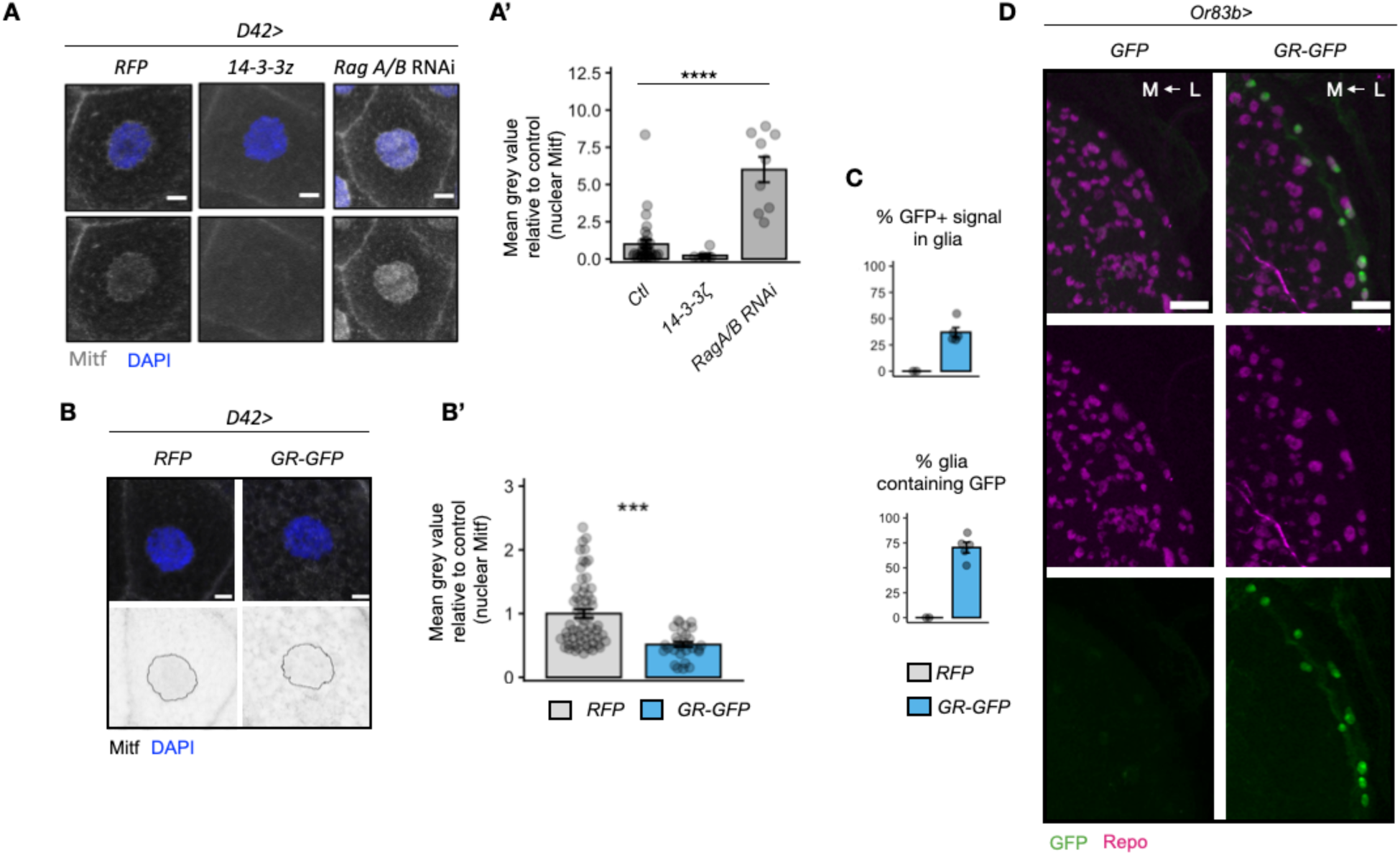
PolyGR-induced pathology. **(A-A’)** Representative images of larval salivary glands expressing control RFP or *14-3-3ζ,* or *RagA/B* RNAi under *D42-GAL4*, which is also active in larval salivary glands, stained to detect endogenous Mitf and DAPI (A; scale bars = 20µm). Quantification of nuclear Mitf fluorescence intensity. One-way ANOVA with Tukey’s multiple comparisons test (A’). **(B-B’)** Representative images of larval salivary glands expressing RFP or 44R GR-GFP under *D42-GAL4*, stained to detect endogenous Mitf and DAPI (B; scale bars = 20µm). Quantification of nuclear Mitf fluorescence intensity. Data are presented as mean ± SEM. Unpaired Welch’s t-tests (B’). **(C)** Quantification of GFP-positive signal in the CB surface glia. Data are presented as mean ± SEM **(D)** Representative images of the adult optic lobe from adult flies expressing GFP or GR-GFP in olfactory sensory neurons with *Or83b-GAL4*, stained to detect GFP and Repo. (C; scale bars = 20µm). GR-GFP is detected predominantly in outer lamina-associated glia (L), with little or no detectable signal in more medially located glia (M).

**Supplementary table 1.** *Drosophila* lines used in this study. Where available, UAS-RNAi lines are accompanied by representative publications validating their use in vivo. VDRC = Vienna drosophila resource center; BL = Bloomington

| Line | Origin |
|---|---|
| UAS-RagA/B RNAi | VDRC 20130 |
| UAS-Mitf RNAi | VDRC 108519 |
| UAS-Vha14 RNAi | VDRC 110160 |
| UAS-Vha100 RNAi | VDRC 108905 |
| UAS-VhaPP1 RNAi | VDRC 47188 |
| UAS-Vha44 RNAi | BL 33884 |
| UAS-VhaSFD RNAi | VDRC 47471 |
| UAS-Cathpesin D RNAi | BL 28978 |
| UAS-Atg1 RNAi | BL 26731 |
| UAS-Atg7 RNAi | BL 27707 |
| UAS-Atg5 RNAi | BL 27551 |
| UAS-Atg8a RNAi | BL 34340 |
| UAS-Atg16 RNAi | BL 34358 |
| UAS-ref(2)P RNAi | BL 36111 |
| UAS-Snap29 RNAi (TRiP) | BL 25862 |
| UAS-Snap29 RNAi (KK) | VDRC 107947 |
| UAS-Sec22 RNAi | BL 34893 |
| UAS-Rab8 RNAi | BL 34373 |
| UAS-Vamp7 RNAi | BL 43543 |
| UAS-Syntaxin17 RNAi | BL 25896 |
| UAS-Ykt6 RNAi | BL 50937 |
| UAS-mCherry RNAi | BL 35785 |
| UAS-mCherry-Atg8a | Gift from Gábor Juhász |
| CathD-mCherry | Gift from Gábor Juhász |
| UAS-14-3-3z | BL85315 |
| UAS-EGFP | BL4531 |
| FRT42D | BL1802 |
| FTD42, Snap29 B6-21 | BL56818 |
| UAS-CTPsyn RNAi | BL31924 |
| UAS-Syntaxin 1A RNAi | BL25811 |
| UAS-Mis12 RNAi | BL38535 |
| UAS-Ndc80 RNAi | BL38260 |
| UAS-Spc105 RNAi | BL35466 |
| UAS-β2 adaptin RNAi | BL28328 |
| UAS-α2 RNAi | BL32866 |

**Supplementary table 2.**
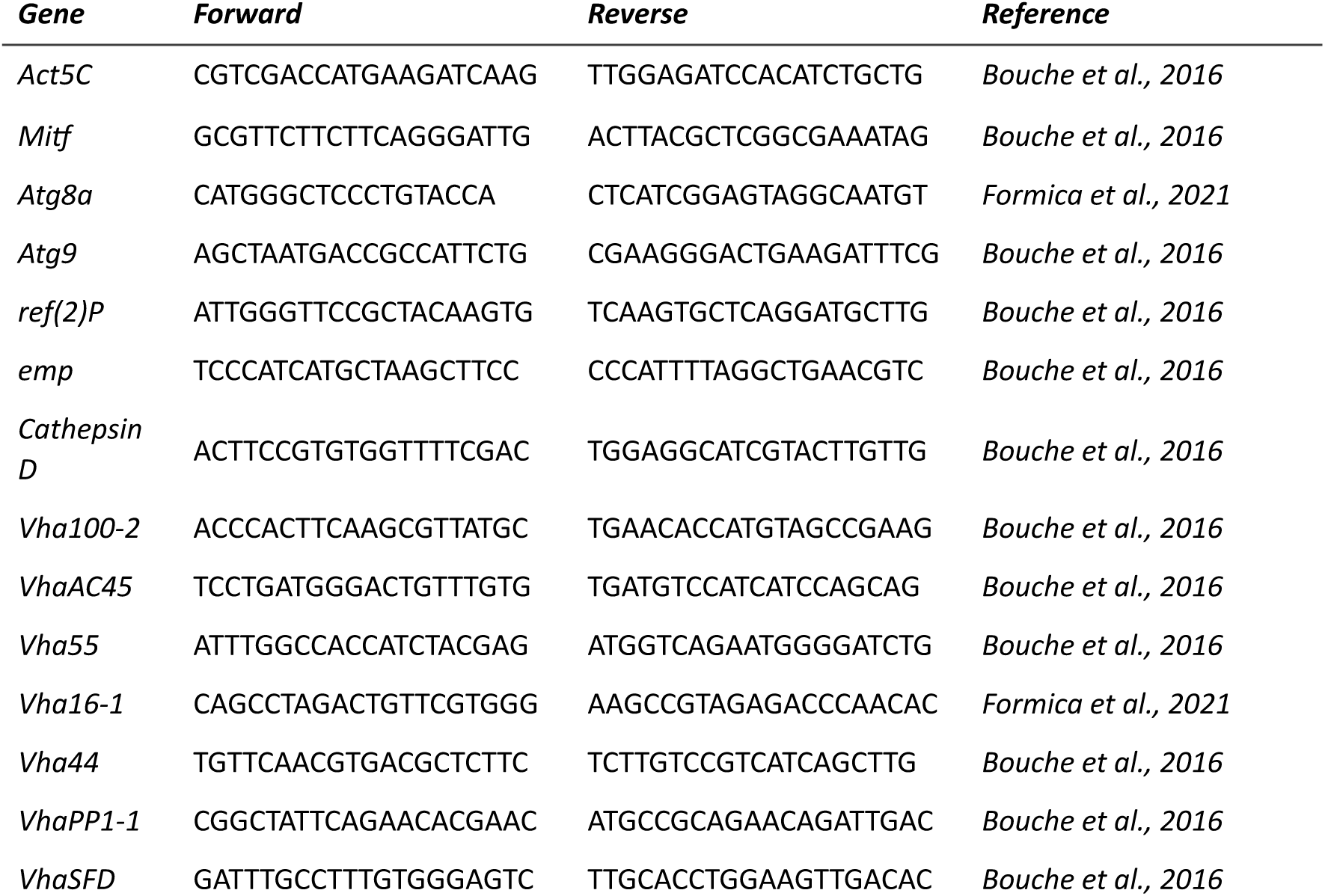
Primers used in this study.

## References

Amick, J., Roczniak-Ferguson, A., & Ferguson, S. M. (2016). C9orf72 binds SMCR8, localizes to lysosomes, and regulates mTORC1 signaling. Molecular Biology of the Cell, 27(20). 10.1091/mbc.E16-01-0003

Andresen, S., Al Outa, A., Formica, M., Enserink, J., & Knævelsrud, H. (2025). Improved detection of lipidated Atg8a by immunoblotting in Drosophila melanogaster cells and tissues enables precise investigation of Atg8a flux and its termination. Autophagy, 21(11), 2497–2517. 10.1080/15548627.2025.2508551

Ash, P. E. A., Bieniek, K. F., Gendron, T. F., Caulfield, T., Lin, W.-L., DeJesus-Hernandez, M., van Blitterswijk, M. M., Jansen-West, K., Paul, J. W., Rademakers, R., Boylan, K. B., Dickson, D. W., & Petrucelli, L. (2013). Unconventional translation of C9ORF72 GGGGCC expansion generates insoluble polypeptides specific to c9FTD/ALS. Neuron, 77(4), 639–646. 10.1016/j.neuron.2013.02.004

Au, W. H., Miller-Fleming, L., Sanchez-Martinez, A., Lee, J. A., Twyning, M. J., Prag, H. A., Raik, L., Allen, S. P., Shaw, P. J., Ferraiuolo, L., Mortiboys, H., & Whitworth, A. J. (2024). Activation of the Keap1/Nrf2 pathway suppresses mitochondrial dysfunction, oxidative stress, and motor phenotypes in C9orf72 ALS/FTD models. Life Science Alliance, 7(9), e202402853. 10.26508/lsa.202402853

Beckers, J., Tharkeshwar, A. K., Fumagalli, L., Contardo, M., Van Schoor, E., Fazal, R., Thal, D. R., Chandran, S., Mancuso, R., Van Den Bosch, L., & Van Damme, P. (2023). A toxic gain-of-function mechanism in C9orf72 ALS impairs the autophagy-lysosome pathway in neurons. Acta Neuropathologica Communications, 11(1). 10.1186/s40478-023-01648-0

Birger, A., Ben-Dor, I., Ottolenghi, M., Turetsky, T., Gil, Y., Sweetat, S., Perez, L., Belzer, V., Casden, N., Steiner, D., Izrael, M., Galun, E., Feldman, E., Behar, O., & Reubinoff, B. (2019). Human iPSC-derived astrocytes from ALS patients with mutated C9ORF72 show increased oxidative stress and neurotoxicity. EBioMedicine, 50, 274–289. 10.1016/j.ebiom.2019.11.026

Boivin, M., Pfister, V., Gaucherot, A., Ruffenach, F., Negroni, L., Sellier, C., & Charlet-Berguerand, N. (2020). Reduced autophagy upon C9ORF72 loss synergizes with dipeptide repeat protein toxicity in G4C2 repeat expansion disorders. The EMBO Journal, 39(4). 10.15252/embj.2018100574

Boll, M. C., Alcaraz-Zubeldia, M., Rios, C., González-Esquivel, D., & Montes, S. (2025). A phase 2, double-blind, placebo-controlled trial of a valproate/lithium combination in ALS patients. Neurología (English Edition), 40(1), 32–40. 10.1016/j.nrleng.2022.07.003

Bonini, N. M., & Fortini, M. E. (2003). Human neurodegenerative disease modeling using Drosophila. Annual Review of Neuroscience, 26, 627–656. 10.1146/annurev.neuro.26.041002.131425

Bouché, V., Perez Espinosa, A., Leone, L., Sardiello, M., Ballabio, A., & Botas, J. (2016). Drosophila Mitf regulates the V-ATPase and the lysosomal-autophagic pathway. Autophagy, 12(3), 484–498. 10.1080/15548627.2015.1134081

Božič, J., Motaln, H., Janež, A. P., Markič, L., Tripathi, P., Yamoah, A., Aronica, E., Lee, Y.-B., Heilig, R., Fischer, R., Thompson, A. J., Goswami, A., & Rogelj, B. (2022). Interactome screening of C9orf72 dipeptide repeats reveals VCP sequestration and functional impairment by poly-GA. Brain, 145(2), 684–699. 10.1093/brain/awab300

Chang, Y. C., Gao, Y., Lee, J. Y., Peng, Y. J., Langen, J., & Chang, K. T. (2024). Identification of secretory autophagy as a mechanism modulating activity-induced synaptic remodeling. Proceedings of the National Academy of Sciences of the United States of America, 121(16), e2315958121. 10.1073/pnas.2315958121

Chang, Y. J., Jeng, U. S., Chiang, Y. L., Hwang, I. S., & Chen, Y. R. (2016). The glycine-alanine dipeptide repeat from C9orf72 hexanucleotide expansions forms toxic amyloids possessing cell-to-cell transmission properties. Journal of Biological Chemistry, 291(10). 10.1074/jbc.M115.694273

Corbier, C., & Sellier, C. (2017). C9ORF72 is a GDP/GTP exchange factor for Rab8 and Rab39 and regulates autophagy. Small GTPases, 8(3). 10.1080/21541248.2016.1212688

Cristofani, R., Crippa, V., Vezzoli, G., Rusmini, P., Galbiati, M., Cicardi, M. E., Meroni, M., Ferrari, V., Tedesco, B., Piccolella, M., Messi, E., Carra, S., & Poletti, A. (2018). The small heat shock protein B8 (HSPB8) efficiently removes aggregating species of dipeptides produced in C9ORF72-related neurodegenerative diseases. Cell Stress & Chaperones, 23(1), 1–12. 10.1007/s12192-017-0806-9

Cunningham, K. M., Maulding, K., Ruan, K., Senturk, M., Grima, J. C., Sung, H., Zuo, Z., Song, H., Gao, J., Dubey, S., Rothstein, J. D., Zhang, K., Bellen, H. J., & Lloyd, T. E. (2020). TFEB/Mitf links impaired nuclear import to autophagolysosomal dysfunction in C9-ALS. eLife, 9, e59419. 10.7554/eLife.59419

Damrau, C., Toshima, N., Tanimura, T., Brembs, B., & Colomb, J. (2018). Octopamine and tyramine contribute separately to the counter-regulatory response to sugar deficit in Drosophila. Frontiers in Systems Neuroscience, 11, 100. 10.3389/fnsys.2017.00100

DeJesus-Hernandez, M., Mackenzie, I. R., Boeve, B. F., Boxer, A. L., Baker, M., Rutherford, N. J., Nicholson, A. M., Finch, N. A., Flynn, H., Adamson, J., Kouri, N., Wojtas, A., Sengdy, P., Hsiung, G.-Y. R., Karydas, A., Seeley, W. W., Josephs, K. A., Coppola, G., Geschwind, D. H., … Rademakers, R. (2011). Expanded GGGGCC hexanucleotide repeat in noncoding region of C9ORF72 causes chromosome 9p-linked FTD and ALS. Neuron, 72(2), 245–256. 10.1016/j.neuron.2011.09.011

Dupont, N., Jiang, S., Pilli, M., Ornatowski, W., Bhattacharya, D., & Deretic, V. (2011). Autophagy-based unconventional secretory pathway for extracellular delivery of IL-1β. The EMBO Journal, 30(23). 10.1038/emboj.2011.398

Ejlerskov, P., Rasmussen, I., Nielsen, T. T., Bergström, A.-L., Tohyama, Y., Jensen, P. H., & Vilhardt, F. (2013). Tubulin polymerization-promoting protein (TPPP/p25α) promotes unconventional secretion of α-synuclein through exophagy by impairing autophagosome-lysosome fusion. Journal of Biological Chemistry, 288(24), 17313–17335. 10.1074/jbc.M112.401174

Ferrari, V., Tedesco, B., Cozzi, M., Pramaggiore, P., Gagliani, M. C., Magdalena, R., Cornaggia, L., Casarotto, E., Chierichetti, M., Mohamed, A., Brodnanovà, M., Milioto, C., Piccolella, M., Galbiati, M., Crippa, V., Provenzani, A., Cortese, K., Rusmini, P., Cristofani, R., & Poletti, A. (2026). VCP modulation ameliorates pathological features in C9orf72 models. Cell Death & Disease, 17, 629. 10.1038/s41419-026-08856-1

Freibaum, B. D., Lu, Y., Lopez-Gonzalez, R., Kim, N. C., Almeida, S., Lee, K. H., Badders, N., Valentine, M., Miller, B. L., Wong, P. C., Petrucelli, L., Kim, H. J., Gao, F. B., & Taylor, J. P. (2015). GGGGCC repeat expansion in C9orf72 compromises nucleocytoplasmic transport. Nature, 525(7567). 10.1038/nature14974

Gendron, T. F., Bieniek, K. F., Zhang, Y.-J., Jansen-West, K., Ash, P. E. A., Caulfield, T., Daughrity, L., Dunmore, J. H., Castanedes-Casey, M., Chew, J., Cosio, D. M., van Blitterswijk, M., Lee, W. C., Rademakers, R., Boylan, K. B., Dickson, D. W., & Petrucelli, L. (2013). Antisense transcripts of the expanded C9ORF72 hexanucleotide repeat form nuclear RNA foci and undergo repeat-associated non-ATG translation in c9FTD/ALS. Acta Neuropathologica, 126(6), 829–844. 10.1007/s00401-013-1192-8

Hartmann, J., Bajaj, T., Otten, J., Klengel, C., Ebert, T., Gellner, A.-K., Junglas, E., Hafner, K., Anderzhanova, E. A., Tang, F., Missig, G., Rexrode, L., Trussell, D. T., Li, K. X., Pöhlmann, M. L., Mackert, S., Geiger, T. M., Heinz, D. E., Lardenoije, R., … Gassen, N. C. (2024). SKA2-regulated hyperactive secretory autophagy drives neuroinflammation-induced neurodegeneration. Nature Communications, 15(1), 2635. 10.1038/s41467-024-46953-x

Hutchings, A.-J., Hambrecht, B., Veh, A., Giridhar, N. J., Zare, A., Angerer, C., Ohnesorge, T., Schenke, M., Selvaraj, B. T., Chandran, S., Sterneckert, J., Petri, S., Seeger, B., Briese, M., Stigloher, C., Bischler, T., Hermann, A., Damme, M., Sendtner, M., & Lüningschrör, P. (2024). Plekhg5 controls the unconventional secretion of Sod1 by presynaptic secretory autophagy. Nature Communications, 15, 8622. 10.1038/s41467-024-52875-5

Juhász, G., Erdi, B., Sass, M., & Neufeld, T.P. (2007). Atg7-dependent autophagy promotes neuronal health, stress tolerance, and longevity but is dispensable for metamorphosis in Drosophila. Genes & Devevelopment, 21(23), 3061–6. 10.1101/gad.1600707

Kang, S., Son, S. M., Baik, S. H., Yang, J., & Mook-Jung, I. (2019). Autophagy-mediated secretory pathway is responsible for both normal and pathological tau in neurons. Journal of Alzheimer’s Disease, 70(3), 667–680. 10.3233/JAD-190180

Khosravi, B., LaClair, K. D., Riemenschneider, H., Zhou, Q., Frottin, F., Mareljic, N., Czuppa, M., Farny, D., Hartmann, H., Michaelsen, M., Arzberger, T., Hartl, F. U., Hipp, M. S., & Edbauer, D. (2020). Cell-to-cell transmission of C9orf72 poly-(Gly-Ala) triggers key features of ALS/FTD. The EMBO Journal, 39(8). 10.15252/embj.2019102811

Kim, J., Yu, Y. S., Choi, Y., Lee, D. H., Han, S., Kwon, J., Noda, T., Ikawa, M., Kim, D., Kim, H., Ballabio, A., Kim, K. I., & Baek, S. H. (2024). USF2 and TFEB compete in regulating lysosomal and autophagy genes. Nature Communications, 15(1), 8334. 10.1038/s41467-024-52600-2

Kimura, T., Jia, J., Kumar, S., Choi, S. W., Gu, Y., Mudd, M., Dupont, N., Jiang, S., Peters, R., Farzam, F., Jain, A., Lidke, K. A., Adams, C. M., Johansen, T., & Deretic, V. (2017). Dedicated SNAREs and specialized TRIM cargo receptors mediate secretory autophagy. The EMBO Journal, 36(1). 10.15252/embj.201695081

Lehmer, C., Oeckl, P., Weishaupt, J. H., Volk, A. E., Diehl-Schmid, J., Schroeter, M. L., Lauer, M., Kornhuber, J., Levin, J., Fassbender, K., Landwehrmeyer, B., Schludi, M. H., Arzberger, T., Kremmer, E., Flatley, A., Feederle, R., Steinacker, P., Weydt, P., Ludolph, A. C., … Otto, M. (2017). Poly-GP in cerebrospinal fluid links C9orf72-associated dipeptide repeat expression to the asymptomatic phase of ALS/FTD. EMBO Molecular Medicine, 9(7), 859–868. 10.15252/emmm.201607486

Leidal, A. M., Huang, H. H., Marsh, T., Solvik, T., Zhang, D., Ye, J., Kai, F., Goldsmith, J., Liu, J. Y., Huang, Y.-H., Monkkonen, T., Vlahakis, A., Huang, E. J., Goodarzi, H., Yu, L., Wiita, A. P., & Debnath, J. (2020). The LC3-conjugation machinery specifies the loading of RNA-binding proteins into extracellular vesicles. Nature Cell Biology, 22(2), 187–199. 10.1038/s41556-019-0450-y

Lin, C.-Y., Wu, H.-E., Weng, E. F.-J., Wu, H.-C., Su, T.-P., & Wang, S.-M. (2024). Fluvoxamine exerts Sigma-1R to rescue autophagy via Pom121-mediated nucleocytoplasmic transport of TFEB. Molecular Neurobiology, 61(8), 5282–5294. 10.1007/s12035-023-03885-9

Ling, S.-C., Polymenidou, M., & Cleveland, D. W. (2013). Converging mechanisms in ALS and FTD: Disrupted RNA and protein homeostasis. Neuron, 79(3), 416–438. 10.1016/j.neuron.2013.07.033

Mackenzie, I. R. A., Frick, P., Grässer, F. A., Gendron, T. F., Petrucelli, L., Cashman, N. R., Edbauer, D., Kremmer, E., Prudlo, J., Troost, D., & Neumann, M. (2015). Quantitative analysis and clinico-pathological correlations of different dipeptide repeat protein pathologies in C9ORF72 mutation carriers. Acta Neuropathologica, 130(6). 10.1007/s00401-015-1476-2

Masrori, P., & Van Damme, P. (2020). Amyotrophic lateral sclerosis: A clinical review. European Journal of Neurology, 27(10), 1918–1929. 10.1111/ene.14393

Mastrodonato, V., Beznoussenko, G., Mironov, A., Ferrari, L., Deflorian, G., & Vaccari, T. (2019). A genetic model of CEDNIK syndrome in zebrafish highlights the role of the SNARE protein Snap29 in neuromotor and epidermal development. Scientific Reports, 9, 1211. 10.1038/s41598-018-37780-4

Mastrodonato, V., Morelli, E., & Vaccari, T. (2018). How to use a multipurpose SNARE: The emerging role of Snap29 in cellular health. Cell Stress, 2(4), 72–81. 10.15698/cst2018.04.130

May, S., Hornburg, D., Schludi, M. H., Arzberger, T., Rentzsch, K., Schwenk, B. M., Grässer, F. A., Mori, K., Kremmer, E., Banzhaf-Strathmann, J., Mann, M., Meissner, F., & Edbauer, D. (2014). C9orf72 FTLD/ALS-associated Gly-Ala dipeptide repeat proteins cause neuronal toxicity and Unc119 sequestration. Acta Neuropathologica, 128(4). 10.1007/s00401-014-1329-4

Mizielinska, S., Grönke, S., Niccoli, T., Ridler, C. E., Clayton, E. L., Devoy, A., Moens, T., Norona, F. E., Woollacott, I. O. C., Pietrzyk, J., Cleverley, K., Nicoll, A. J., Pickering-Brown, S., Dols, J., Cabecinha, M., Hendrich, O., Fratta, P., Fisher, E. M. C., Partridge, L., & Isaacs, A. M. (2014). C9orf72 repeat expansions cause neurodegeneration in Drosophilathrough arginine-rich proteins. Science, 345(6201), 1192–1194. 10.1126/science.1256800

Morelli, E., Ginefra, P., Mastrodonato, V., Beznoussenko, G. V., Rusten, T. E., Bilder, D., Stenmark, H., Mironov, A. A., & Vaccari, T. (2014). Multiple functions of the SNARE protein Snap29 in autophagy, endocytic, and exocytic trafficking during epithelial formation in Drosophila. Autophagy, 10(12), 2251–2268. 10.4161/15548627.2014.981913

Morelli, E., Speranza, E. A., Pellegrino, E., Beznoussenko, G. V., Carminati, F., Garré, M., Mironov, A. A., Onorati, M., & Vaccari, T. (2021). Activity of the SNARE protein SNAP29 at the endoplasmic reticulum and Golgi apparatus. Frontiers in Cell and Developmental Biology, 9, 637565. 10.3389/fcell.2021.637565

Mori, K., Weng, S.-M., Arzberger, T., May, S., Rentzsch, K., Kremmer, E., Schmid, B., Kretzschmar, H. A., Cruts, M., Van Broeckhoven, C., Haass, C., & Edbauer, D. (2013). The C9orf72 GGGGCC repeat is translated into aggregating dipeptide-repeat proteins in FTLD/ALS. Science, 339(6125), 1335–1338. 10.1126/science.1232927

Morón-Oset, J., Supèr, T., Esser, J., Isaacs, A. M., Grönke, S., & Partridge, L. (2019). Glycine-alanine dipeptide repeats spread rapidly in a repeat length- and age-dependent manner in the fly brain. Acta Neuropathologica Communications, 7, 209. 10.1186/s40478-019-0860-x

Nakamura, Y., Sawai, T., Kakiuchi, K., & Arawaka, S. (2024). Neuronal activity promotes secretory autophagy for the extracellular release of α-synuclein. Journal of Biological Chemistry, 300(7), 107419. 10.1016/j.jbc.2024.107419

Nezis, I. P., Shravage, B. V., Sagona, A. P., Lamark, T., Bjørkøy, G., Johansen, T., Rusten, T. E., Brech, A., Baehrecke, E. H., & Stenmark, H. (2010). Autophagic degradation of dBruce controls DNA fragmentation in nurse cells during late Drosophila melanogaster oogenesis. Journal of Cell Biology, 190(4), 523–531. 10.1083/jcb.201002035

Nilsson, P., Loganathan, K., Sekiguchi, M., Matsuba, Y., Hui, K., Tsubuki, S., Tanaka, M., Iwata, N., Saito, T., & Saido, T. C. (2013). Aβ secretion and plaque formation depend on autophagy. Cell Reports, 5(1), 61–69. 10.1016/j.celrep.2013.08.042

O’Rourke, J. G., Bogdanik, L., Yáñez, A., Lall, D., Wolf, A. J., Muhammad, A. K. M. G., Ho, R., Carmona, S., Vit, J. P., Zarrow, J., Kim, K. J., Bell, S., Harms, M. B., Miller, T. M., Dangler, C. A., Underhill, D. M., Goodridge, H. S., Lutz, C. M., & Baloh, R. H. (2016). C9orf72 is required for proper macrophage and microglial function in mice. Science, 351(6279). 10.1126/science.aaf1064

Pajares, M., Jiménez-Moreno, N., García-Yagüe, Á. J., Escoll, M., de Ceballos, M. L., Van Leuven, F., Rábano, A., Yamamoto, M., Rojo, A. I., & Cuadrado, A. (2016). Transcription factor NFE2L2/NRF2 is a regulator of macroautophagy genes. Autophagy, 12(10), 1902–1916. 10.1080/15548627.2016.1208889

Piepers, S., Veldink, J. H., De Jong, S. W., Van Der Tweel, I., Van Der Pol, W.-L., Uijtendaal, E. V., Schelhaas, H. J., Scheffer, H., De Visser, M., De Jong, J. M. B. V., Wokke, J. H. J., Groeneveld, G. J., & Van Den Berg, L. H. (2009). Randomized sequential trial of valproic acid in amyotrophic lateral sclerosis. Annals of Neurology, 66(2), 227–234. 10.1002/ana.21620

Platt, F. M., d’Azzo, A., Davidson, B. L., Neufeld, E. F., & Tifft, C. J. (2018). Lysosomal storage diseases. Nature Reviews Disease Primers, 4, 27. 10.1038/s41572-018-0025-4

Ramos-Campoy, O., Ávila-Polo, R., Grau-Rivera, O., Antonell, A., Clarimón, J., Rojas-García, R., Charif, S., Santiago-Valera, V., Hernandez, I., Aguilar, M., Almenar, C., Lopez-Villegas, D., Bajo, L., Pastor, P., Van der Zee, J., Lladó, A., Sanchez-Valle, R., & Gelpi, E. (2018). Systematic screening of ubiquitin/p62 aggregates in cerebellar cortex expands the neuropathological phenotype of the C9orf72 expansion mutation. Journal of Neuropathology & Experimental Neurology, 77(8), 703–709. 10.1093/jnen/nly047

Renton, A. E., Majounie, E., Waite, A., Simón-Sánchez, J., Rollinson, S., Gibbs, J. R., Schymick, J. C., Laaksovirta, H., van Swieten, J. C., Myllykangas, L., Kalimo, H., Paetau, A., Abramzon, Y., Remes, A. M., Kaganovich, A., Scholz, S. W., Duckworth, J., Ding, J., Harmer, D. W., … Traynor, B. J. (2011). A hexanucleotide repeat expansion in C9ORF72 is the cause of chromosome 9p21-linked ALS-FTD. Neuron, 72(2), 257–268. 10.1016/j.neuron.2011.09.010

Riemslagh, F. W., Verhagen, R. F. M., van der Toorn, E. C., Smits, D. J., Quint, W. H., van der Linde, H. C., van Ham, T. J., & Willemsen, R. (2021). Reduction of oxidative stress suppresses poly-GR-mediated toxicity in zebrafish embryos. Disease Models & Mechanisms, 14(11), dmm049092. 10.1242/dmm.049092

Sawai, T., Nakamura, Y., & Arawaka, S. (2025). Secretory autophagy mediates lysosomal and autophagic degradation for α-synuclein proteostasis. Journal of Biological Chemistry, 301(8), 110474. 10.1016/j.jbc.2025.110474

Schiller, S. A., Seebode, C., Wieser, G. L., Goebbels, S., Möbius, W., Horowitz, M., Sarig, O., Sprecher, E., & Emmert, S. (2016). Establishment of two mouse models for CEDNIK syndrome reveals the pivotal role of SNAP29 in epidermal differentiation. Journal of Investigative Dermatology, 136(3), 672–679. 10.1016/j.jid.2015.12.020

Smeele, P. H., & Vaccari, T. (2023). Snapshots from within the cell: Novel trafficking and non-trafficking functions of Snap29 during tissue morphogenesis. Seminars in Cell & Developmental Biology, 133, 42–52. 10.1016/j.semcdb.2022.02.024

Solvik, T. A., Nguyen, T. A., Lin, Y.-H. T., Marsh, T., Huang, E. J., Wiita, A. P., Debnath, J., & Leidal, A. M. (2022). Secretory autophagy maintains proteostasis upon lysosome inhibition. Journal of Cell Biology, 221(6), e202110151. 10.1083/jcb.202110151

Su, Z., Zhang, Y., Gendron, T. F., Bauer, P. O., Chew, J., Yang, W.-Y., Fostvedt, E., Jansen-West, K., Belzil, V. V., Desaro, P., Johnston, A., Overstreet, K., Oh, S.-Y., Todd, P. K., Berry, J. D., Cudkowicz, M. E., Boeve, B. F., Dickson, D., Floeter, M. K., … Disney, M. D. (2014). Discovery of a biomarker and lead small molecules to target r(GGGGCC)-associated defects in c9FTD/ALS. Neuron, 83(5), 1043–1050. 10.1016/j.neuron.2014.07.041

Sullivan, P. M., Zhou, X., Robins, A. M., Paushter, D. H., Kim, D., Smolka, M. B., & Hu, F. (2016). The ALS/FTLD-associated protein C9orf72 associates with SMCR8 and WDR41 to regulate the autophagy-lysosome pathway. Acta Neuropathologica Communications, 4(1). 10.1186/s40478-016-0324-5

Sung, H., & Lloyd, T. E. (2023). Disrupted endoplasmic reticulum-mediated autophagosomal biogenesis in a Drosophila model of C9-ALS-FTD. Autophagy. 10.1080/15548627.2023.2249750

Takáts, S., Nagy, P., Varga, Á., Pircs, K., Kárpáti, M., Varga, K., Kovács, A. L., Hegedűs, K., & Juhász, G. (2013). Autophagosomal Syntaxin17-dependent lysosomal degradation maintains neuronal function in Drosophila. Journal of Cell Biology, 201(4), 531–539. 10.1083/jcb.201211160

Tognon, E., Kobia, F., Busi, I., Fumagalli, A., De Masi, F., & Vaccari, T. (2016). Control of lysosomal biogenesis and Notch-dependent tissue patterning by components of the TFEB-V-ATPase axis in Drosophila melanogaster. Autophagy, 12(3), 499–514. 10.1080/15548627.2015.1134080

Wang, S. M., Wu, H. E., Yasui, Y., Geva, M., Hayden, M., Maurice, T., Cozzolino, M., & Su, T. P. (2023). Nucleoporin POM121 signals TFEB-mediated autophagy via activation of SIGMAR1/sigma-1 receptor chaperone by pridopidine. Autophagy, 19(1). 10.1080/15548627.2022.2063003

Webster, C. P., Smith, E. F., Bauer, C. S., Moller, A., Hautbergue, G. M., Ferraiuolo, L., Myszczynska, M. A., Higginbottom, A., Walsh, M. J., Whitworth, A. J., Kaspar, B. K., Meyer, K., Shaw, P. J., Grierson, A. J., & De Vos, K. J. (2016). The C9orf72 protein interacts with Rab1a and the ULK1 complex to regulate initiation of autophagy. The EMBO Journal, 35(15). 10.15252/embj.201694401

Westergard, T., Jensen, B. K., Wen, X., Cai, J., Kropf, E., Iacovitti, L., Pasinelli, P., & Trotti, D. (2016). Cell-to-cell transmission of dipeptide repeat proteins linked to C9orf72-ALS/FTD. Cell Reports, 17(3). 10.1016/j.celrep.2016.09.032

Xu, D., Vincent, A., González-Gutiérrez, A., Aleyakpo, B., Anoar, S., Giblin, A., Atilano, M. L., Adams, M., Shen, D., Thoeng, A., Tsintzas, E., Maeland, M., Isaacs, A. M., Sierralta, J., & Niccoli, T. (2023). A monocarboxylate transporter rescues frontotemporal dementia and Alzheimer’s disease models. PLOS Genetics, 19(9), e1010893. 10.1371/journal.pgen.1010893

Xu, J., Yang, K. C., Go, N. E., Colborne, S., Ho, C. J., Hosseini-Beheshti, E., Lystad, A. H., Simonsen, A., Guns, E. T., Morin, G. B., & Gorski, S. M. (2022). Chloroquine treatment induces secretion of autophagy-related proteins and inclusion of Atg8-family proteins in distinct extracellular vesicle populations. Autophagy, 18(11), 2547–2560. 10.1080/15548627.2022.2039535

Yamakawa, M., Ito, D., Honda, T., Kubo, K.-I., Noda, M., Nakajima, K., & Suzuki, N. (2015). Characterization of the dipeptide repeat protein in the molecular pathogenesis of c9FTD/ALS. Human Molecular Genetics, 24(6), 1630–1645. 10.1093/hmg/ddu576

Yang, M., Chen, L., Swaminathan, K., Herrlinger, S., Lai, F., Shiekhattar, R., & Chen, J. F. (2016). A C9ORF72/SMCR8-containing complex regulates ULK1 and plays a dual role in autophagy. Science Advances, 2(9). 10.1126/sciadv.1601167

Yi, J., Zhang, L., Tang, B., Han, W., Zhou, Y., Chen, Z., Jia, D., & Jiang, H. (2013). Sodium valproate alleviates neurodegeneration in SCA3/MJD via suppressing apoptosis and rescuing the hypoacetylation levels of histone H3 and H4. PLOS ONE, 8(1), e54792. 10.1371/journal.pone.0054792

Zhang, K., Donnelly, C. J., Haeusler, A. R., Grima, J. C., Machamer, J. B., Steinwald, P., Daley, E. L., Miller, S. J., Cunningham, K. M., Vidensky, S., Gupta, S., Thomas, M. A., Hong, I., Chiu, S. L., Huganir, R. L., Ostrow, L. W., Matunis, M. J., Wang, J., Sattler, R., … Rothstein, J. D. (2015). The C9orf72 repeat expansion disrupts nucleocytoplasmic transport. Nature, 525(7567). 10.1038/nature14973

Zhang, Y.-J., Guo, L., Gonzales, P. K., Gendron, T. F., Wu, Y., Jansen-West, K., O’Raw, A. D., Pickles, S. R., Prudencio, M., Carlomagno, Y., Gachechiladze, M. A., Ludwig, C., Tian, R., Chew, J., DeTure, M., Lin, W.-L., Tong, J., Daughrity, L. M., Yue, M., … Petrucelli, L. (2019). Heterochromatin anomalies and double-stranded RNA accumulation underlie C9orf72 poly(PR) toxicity. Science, 363(6428), eaav2606. 10.1126/science.aav2606

Zhou, Q., Lehmer, C., Michaelsen, M., Mori, K., Alterauge, D., Baumjohann, D., Schludi, M. H., Greiling, J., Farny, D., Flatley, A., Feederle, R., May, S., Schreiber, F., Arzberger, T., Kuhm, C., Klopstock, T., Hermann, A., Haass, C., & Edbauer, D. (2017). Antibodies inhibit transmission and aggregation of C9orf72 poly-GA dipeptide repeat proteins. EMBO Molecular Medicine, 9(5). 10.15252/emmm.201607054

Zirin, J., Nieuwenhuis, J., & Perrimon, N. (2013). Role of autophagy in glycogen breakdown and its relevance to chloroquine myopathy. PLOS Biology, 11(11), e1001708. 10.1371/journal.pbio.1001708

Zu, T., Liu, Y., Bañez-Coronel, M., Reid, T., Pletnikova, O., Lewis, J., Miller, T. M., Harms, M. B., Falchook, A. E., Subramony, S. H., Ostrow, L. W., Rothstein, J. D., Troncoso, J. C., & Ranum, L. P. W. (2013). RAN proteins and RNA foci from antisense transcripts in C9ORF72 ALS and frontotemporal dementia. Proceedings of the National Academy of Sciences of the United States of America, 110(51). 10.1073/pnas.1315438110

